# An improved Erk biosensor reveals oscillatory Erk dynamics driven by mitotic erasure during early development

**DOI:** 10.1101/2022.11.03.515001

**Authors:** Scott G. Wilcockson, Luca Guglielmi, Pablo Araguas Rodriguez, Marc Amoyel, Caroline S. Hill

## Abstract

Erk signaling dynamics elicit distinct cellular responses in a variety of contexts. The early zebrafish embryo is an ideal model to explore the role of Erk signaling dynamics *in vivo*, as a gradient of activated diphosphorylated Erk (P-Erk) is induced by Fgf signaling at the blastula embryonic margin. Here we describe an improved Erk-specific biosensor which we term modified Erk Kinase Translocation Reporter (modErk-KTR). We demonstrate the utility of this biosensor *in vitro* and in developing zebrafish and *Drosophila* embryos. Moreover, we show that Fgf/Erk signaling is dynamic and coupled to tissue growth during both early zebrafish and *Drosophila* development. Signaling is rapidly extinguished just prior to mitosis, which we refer to as mitotic erasure, inducing periods of inactivity, thus providing a source of heterogeneity in an asynchronously dividing tissue. Our modified reporter and transgenic lines represent an important resource for interrogating the role of Erk signaling dynamics *in vivo*.

## Introduction

Embryonic development requires the coordinated communication, proliferation, and movement of cells on a grand scale. The highly conserved extracellular signal regulated kinase (Erk) is a key node connecting these essential processes, as well as playing a critical role in coordinating cell fate specification ^1^. Understanding the regulation and output of Erk signaling is therefore crucial for understanding its role in development as well as in adult homeostasis and disease. Thus, the development of sensitive methods of visualizing signaling Erk dynamics *in vivo* is essential.

Functioning downstream of receptor tyrosine kinase (RTK) receptors, Erk signaling can elicit different cellular responses depending on context and upstream ligand-receptor combinations that can be associated with distinct signaling dynamics. For example, treatment of rat PC-12 cells with EGF (epidermal growth factor) induces transient signaling which promotes proliferation, while NGF (nerve growth factor) induces sustained signaling that promotes differentiation ^2,3^. Introducing regular pulses of EGF, rather than sustained addition, is sufficient to convert EGF to a pro-differentiation signal ^4^. Similarly, while short-term sustained Erk signaling (~30 min) promotes neural fate in the *Drosophila* blastoderm, long-term sustained (≥60 min) or frequent pulses of Erk activity promote endodermal fate ^5,6^. It is therefore proposed that information is encoded within Erk dynamics through the accumulative dose of Erk activity. The advent of Erk biosensors is now enabling the interrogation of Erk dynamics in vivo, and recent work has similarly suggested a role for sustained versus pulsatile signaling in mouse ESC differentiation ^7,8^. This highlights the importance of elucidating the role of Erk signaling dynamics in the regulation of cellular identity and behaviour.

The zebrafish embryo presents an ideal model system to monitor signaling dynamics *in vivo* and previous studies have successfully employed Erk biosensors in the study of wounding and vasculogenesis ^9–11^. During early zebrafish development, the patterns and roles of Fibroblast growth factor Fgf/Erk signaling are well characterised, as successive rounds of signaling pattern first the dorsoventral (DV) axis and then the anteroposterior (AP) axis ^12,13^. Between 3.3 and 3.6 hours post fertilization (hpf) (mid-blastula stage), a discrete domain of *fgf8a*, *fgf3* and *fgf24* expression and corresponding P-Erk gradient is induced by Nodal signaling in the presumptive dorsal organizer ^14,15^. Between 4.3 and 5.3 hpf, Nodal signaling induces expression of fgf3/8a in the marginal-most cells to drive long-range Erk signaling around the embryonic margin ^16,17^. Together, overlapping short-range Nodal and long-range Fgf/Erk signaling induce and pattern the mesodermal and endodermal lineages ^18^. Importantly, snap-shot views of development show a highly heterogeneous pattern of Erk activity, as read out by levels of diphosphorylated Erk (P-Erk) ^19^. This pattern suggests heterogeneity in either the single cell response to Fgf signaling or in Erk signaling dynamics over time.

Here we report the generation of a highly specific and sensitive reporter of Erk signaling, through the modification of the Erk-Kinase Translocation Reporter (KTR) ^20^ that abolishes the reporter’s responses to Cdk1. We hereafter refer to the biosensor as modErk-KTR. These KTR reporters use site-specific phosphorylation by the target kinase to regulate nucleocytoplasmic shuttling of a fluorescent protein. We demonstrate the highly specific sensitivity of the modErk-KTR to Erk signaling in zebrafish embryonic tissues, as well as in *Drosophila* embryonic and larval tissues. Furthermore, we monitor the growth and collapse, following inhibition of signaling, of the Fgf/Erk signaling gradient in the zebrafish blastula. We identify oscillations in Erk signaling associated with mitosis, a process we refer to as mitotic erasure, in both zebrafish and *Drosophila* embryos. This introduces periods of Erk inactivity and couples signaling dynamics to tissue growth, thus providing a source of signaling heterogeneity in an asynchronously dividing tissue.

## Design

We observe off-target Erk-KTR activity during early development that we demonstrate is due to Cdk1 activity. Indeed studies with other Erk biosensors reported similar findings that Cdk1 activity can influence reporter readouts ^21,22^. A recent study addressed this issue with the EKAREV FRET sensor by changing the Erk phosphorylation motifs to remove key lysines that mediate Cdk1 recognition ^22^. The phosphorylation sites of the Erk-KTR are similarly surrounded by lysines, but these residues are essential for the function of the nuclear localisation sequence (NLS) ^20^. It is therefore not possible to modify the Erk-KTR phosphorylation site to reduce Cdk1 interaction. Instead, we have modified a putative cyclin-docking site found within the ELK1-derived Erk docking domain to reduce cyclin–Cdk1 binding and hence substantially improve Erk-specificity of the biosensor.

## Results

### Erk-KTR displays Mek/Erk-independent off-target activity in early zebrafish embryos

To monitor Erk activity *in vivo*, we used a previously developed transgenic zebrafish line (*ubiP*:Erk-KTR-Clover) where Erk-KTR is ubiquitously expressed ^9^. The Erk-KTR consists of an N-terminal Erk-docking domain (derived from human ELK1), an NLS containing Erk consensus phosphorylation sites (TP), a nuclear export sequence (NES) and a green fluorescent protein (Clover) (Figure 1A) ^20^. In the absence of activated P-Erk the NLS is dominant, and the reporter concentrates in the nucleus (Figure 1A, B). Upon Erk activation, phosphorylation of the NLS inhibits its function and the KTR shuttles to the cytoplasm. By measuring the cytoplasmic-to-nuclear (C/N) fluorescence ratio of the reporter, it is possible to measure the real-time levels of Erk activity ^20^.

**Figure 1.**
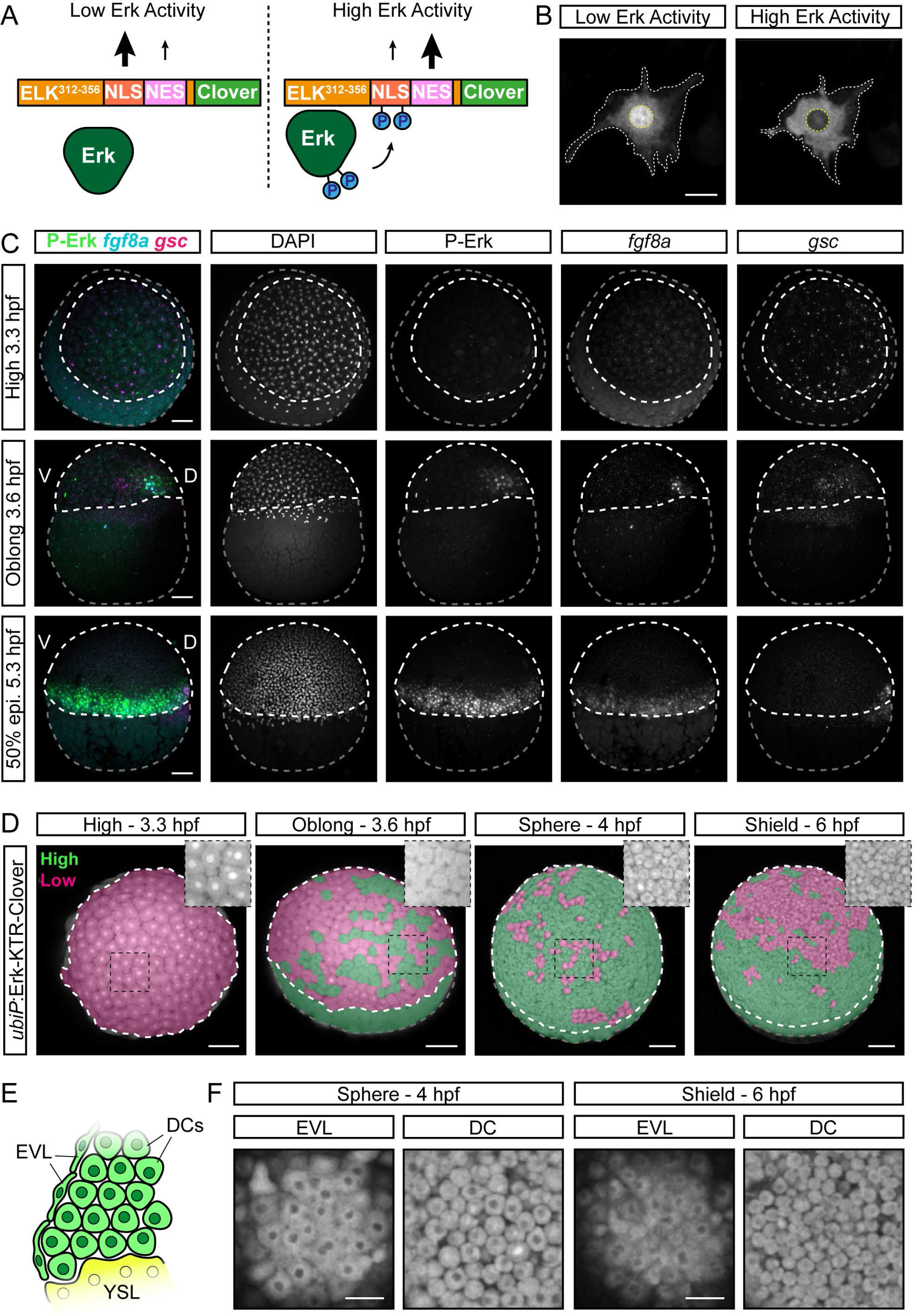
Off-target Erk-KTR activity in the early zebrafish embryo. (A) Schematic of the Erk-KTR construct showing the N-terminal Erk-docking domain derived from ELK1, a nuclear localization sequence (NLS) containing Erk-consensus phosphorylation sites, a nuclear exit sequence (NES) and a C-terminal fluorescent protein, Clover. (B) Live images of an NIH-3T3 cell transfected with *ubiP*:Erk-KTR-Clover construct. The cytoplasmic-to-nuclear ratio of the Erk-KTR fluorescence provides a live readout of relative Erk activity levels. White dashed line, cytoplasm; yellow dashed line, nucleus. (C) Combined immunofluorescence and RNAscope showing a time-course of diphosphorylated Erk (P-Erk) and *fgf8a* expression relative to the dorsal organizer marked by goosecoid (gsc) expression. Embryos are shown from an animal view (3.3 hpf) or lateral view (3.6–5.3 hpf). White dashed line, embryo proper; grey dashed line, yolk; 50% epi., 50% epiboly. (D) Stills of live *ubiP*:Erk-KTR-Clover embryos showing a time course of reporter activity. Embryos are false colored to indicate Erk-KTR activity as readout by the KTR reporter in a binary manner; green shows high Erk-KTR activity and magenta shows low Erk-KTR activity. Embryos are shown from an animal–lateral view. Insets show a magnified view of the region within the black box without false coloring. (E) Schematic of a cross-section of the embryonic margin showing the relative position of the deep cells (DCs), the enveloping layer (EVL) and the yolk syncytial layer (YSL). (F) Single z-slices showing Erk-KTR activity in the EVL and DCs from the indicated embryos in (D). Scalebars, 25 µm (B), 50 µm (F) or 100 µm (C, D).

First, we established whether Erk-KTR reports the known patterns of Erk activity in the zebrafish blastula. P-Erk is first seen at 3.6 hpf in a small dorsal domain colocalizing with gsc and *fgf8a* expression (Figure 1C). By 5.3 hpf, P-Erk is detected throughout the embryonic margin in both deep cells (DCs) and the enveloping layer (EVL) ^16^. To aid the visualization of the Erk-KTR readout, we false-colored embryos in a binary manner to show enrichment in the cytoplasm (green; C:N > 1) or nucleus (magenta; C:N ≤ 1), which indicate high and low Erk activity, respectively (Figure 1D). At 3.3 hpf, the reporter shows strong nuclear localization throughout the blastoderm indicating no Erk activity (Figure 1D). However, from 3.6 hpf we observed sporadic nuclear exclusion throughout the blastoderm, and most cells showed uniform nuclear exclusion by 4.0 hpf, including both DCs and the EVL (Figure 1E, F). By 6.0 hpf, nuclear exclusion of the reporter becomes restricted to the margin, but it was still observed beyond the Fgf/Erk signaling domain (Figure 1D and Video S1). To test whether Erk-KTR localization was Erk-dependent, we measured C/N ratios at both the margin (high Fgf/Erk) and animal pole (no Fgf/Erk; Figure 2A) following treatment from 4.0 to 5.0 hpf with an inhibitor of Mek (mitogen-activated protein kinase kinase), the upstream activator of Erk (10 µM PD-0325901; MEKi). This caused a small but significant decrease in C/N ratios at both the margin and animal pole in comparison with vehicle controls (Figure 2A–C). However, Erk-KTR remained enriched in the cytoplasm in treated embryos, indicating significant Mek/Erk-independent nucleocytoplasmic shuttling in the zebrafish blastula.

**Figure 2.**
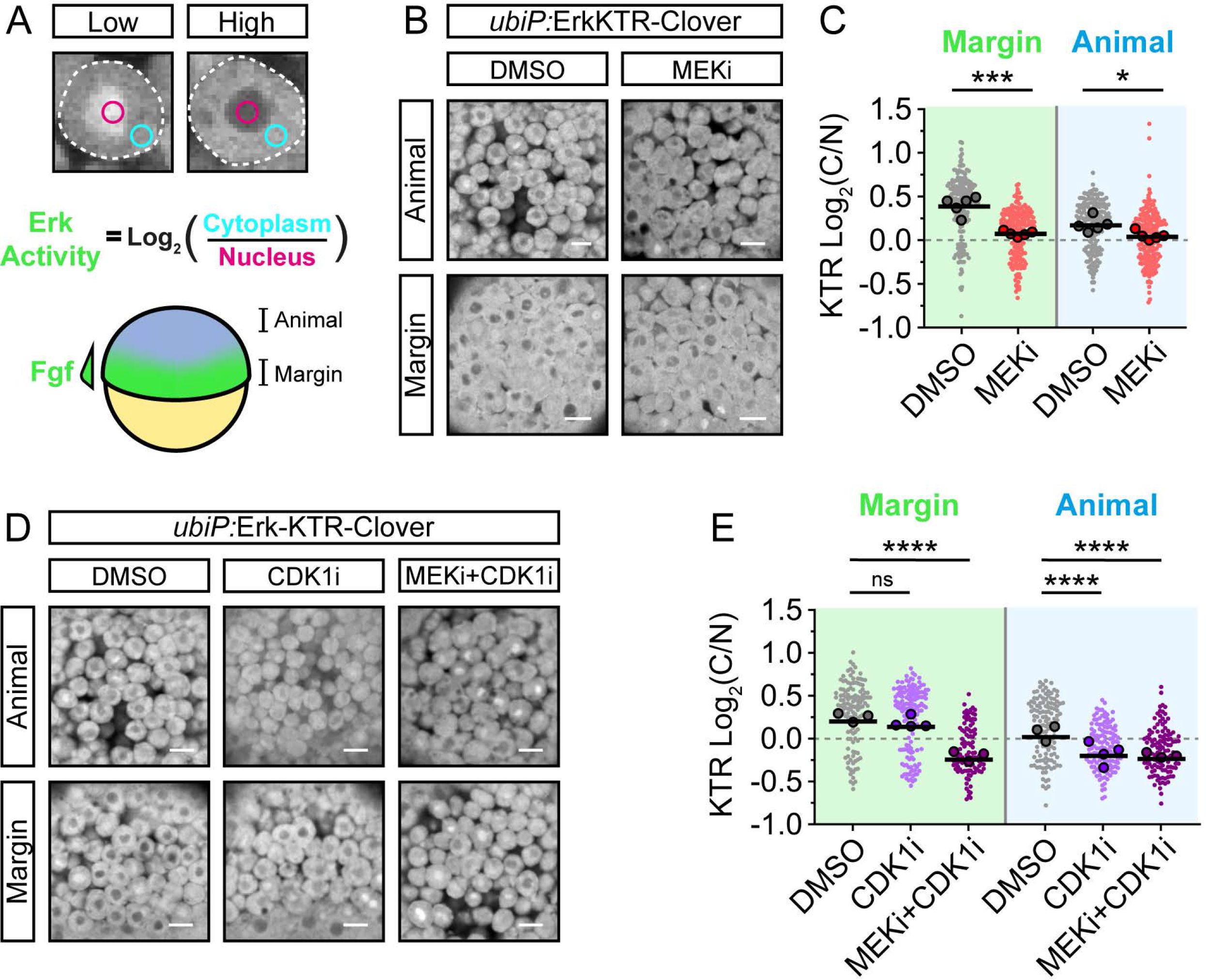
Erk-KTR reports on Erk and Cdk1 activity in early zebrafish embryos. (A) Illustration of the method used to report Erk-KTR activity in early zebrafish embryos (schematized below) by measuring mean fluorescence intensity in a region of the nucleus (magenta) and cytoplasm (cyan). The margin of the embryo exhibits high Fgf signaling, while the animal pole does not. (B) Live imaging of *ubiP*:Erk-KTR-Clover transgenic embryos at either the margin or animally, as indicated in (A), following treatment with DMSO (control) or 10 µM PD-0325901 (MEKi) for an hour from 4.0 hpf. (C) Quantification of Erk-KTR activity in (B) at the margin (p = 0.0002) and animally (p = 0.0207). n = 178–209 cells per condition from 5 embryos. Shown are the single cell readouts of Erk-KTR activity overlayed with the per embryo averages and the overall mean. (D) as in (B) but following treatment with DMSO (control), 20 µM RO-3306 (CDK1i), or both 10 µM PD-0325901 and 20 µM RO-3306 (MEKi+CDK1i) for 1 hr from 4.0 hpf. (E) Quantification of Erk-KTR activity in (B) as in (C) for CDK1i (margin p = 0.3131; animal p < 0.0001) or both MEKi and CDK1i (margin p < 0.0001; animal p < 0.0001). n = 107–146 cells per condition from 3 (DMSO and MEKi+CDK1i) or 4 embryos (CDKi) per condition. Statistical tests were Student t-test (C) or one-way ANOVA with Šidák's multiple comparisons test (E). Scale bars, 20 µm; ****, p <0.0001; ns, not significant.

Previous studies have used the Erk-KTR to monitor signaling dynamics at later stages of development (≥ 24 hpf) ^9,11^ and we observed that by 6.0 hpf the reporter more accurately reflected the expected pattern of Erk signaling (Figure 1D). This suggests that the Mek/Erk-independent shuttling might be driven by some phenomenon occurring during early development. We noted that the onset of reporter mislocalization correlated well with cell cycle remodeling at the midblastula transition ^23,24^. This is driven by lengthening of the cell cycle, which is influenced by changes in Cdc25-Cdk1 activity in zebrafish ^23^. We therefore asked whether Erk-KTR localization was influenced by cell cycle inputs by treating cells with a CDK1-specific inhibitor (20 µM RO-3306; CDK1i). Treatment had no effect on Erk-KTR C/N ratios at the margin, however, there was a significant reduction in C/N ratios animally (Figure 2D, E). Treatment with both MEKi and CDK1i resulted in maximal nuclear localization at the margin. Together, these data show that the localization of Erk-KTR is controlled by a combination of Erk and Cdk1. We propose that the rapid, short cell cycles (every 15–30 min) of early development emphasize this due to an increased frequency of high Cdk1 activity, masking the true pattern of Erk signaling.

### modERK-KTR: a modified Erk-KTR with improved specificity

To enable the monitoring of Fgf/Erk signaling dynamics *in vivo*, we generated an improved Erk biosensor devoid of Cdk1 responsiveness (modErk-KTR). To achieve this, we introduced an R>A substitution within a putative cyclin-docking site (RxLxΦ, where Φ is a hydrophobic residue) in the Erk-docking domain (Figure 3A) ^25^. This would be predicted to significantly reduce cyclin-substrate binding ^26,27^, but may also weakly reduce Erk binding ^28,29^. We therefore introduced another Erk docking site (FQFP) at the C-terminus of the ELK1 fragment (Figure 3A) ^29^.

**Figure 3.**
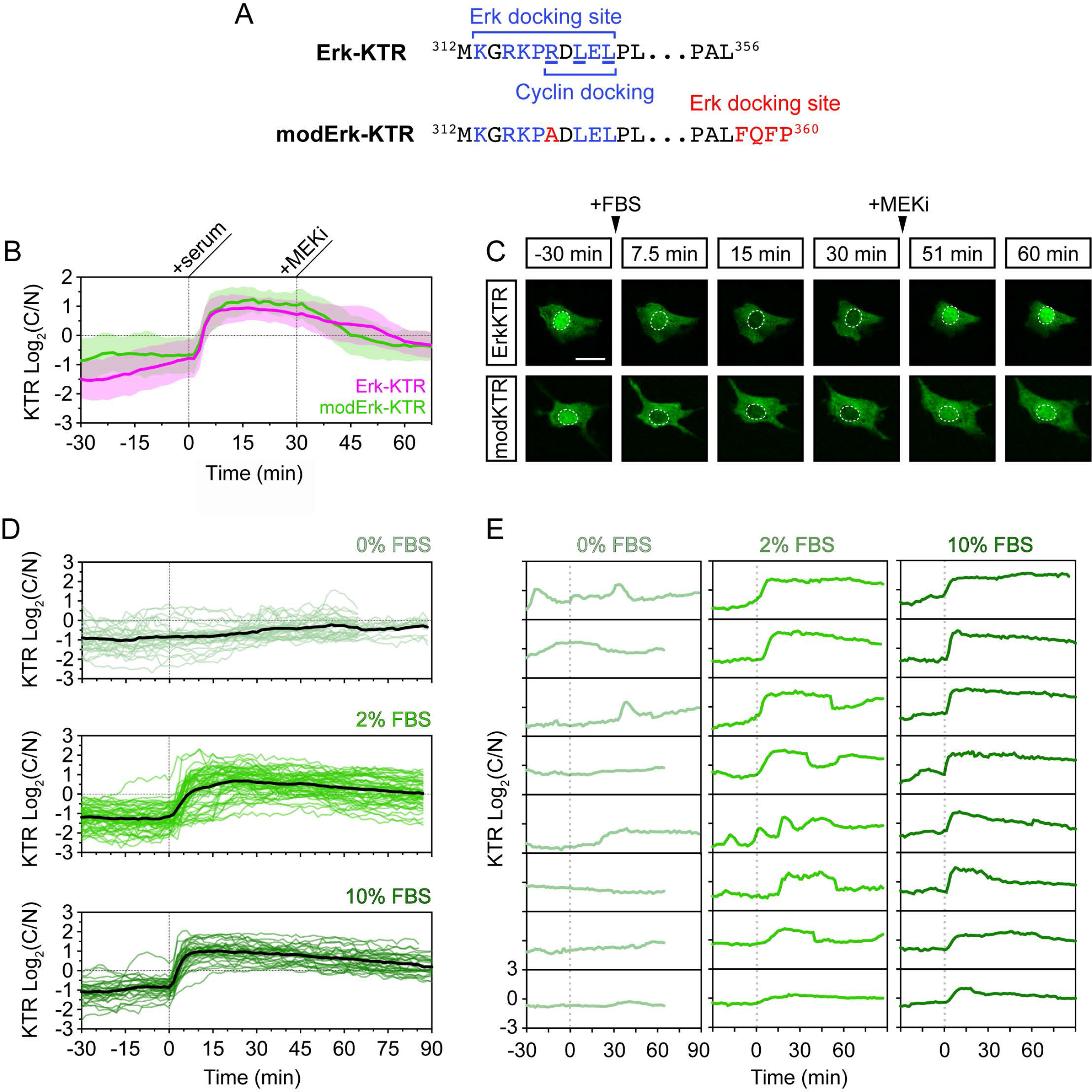
A modified Erk-KTR reports ERK activity in mouse embryonic fibroblasts. (A) Amino acid sequences of the Erk-docking domain of Erk-KTR and modified Erk-KTR highlighting the modifications (red) made to reduce off-target reporter activity, including the R>A substitution within the Erk docking site and the addition of an FQFP Erk-docking site between the ELK fragment and the NLS. (B) Quantification of ERK activity in NIH-3T3 mouse embryonic fibroblasts (n = 58 cells each, mean ± SD). Cells were serum-starved overnight and ERK was induced by the addition of 10% fetal bovine serum (FBS). ERK activity was inhibited after 30 min with 10 µM PD-0325901 (MEKi). (C) Representative images of reporter activity in (B). (D) Quantification of ERK activity in NIH-3T3 cells after overnight serum-starvation, followed by the addition of different concentrations of FBS. Individual cell traces and the mean (black line) are shown for 0% FBS (n = 32 cells), 2% FBS (n = 57 cells) and 10% FBS (n = 30 cells). (E) Individual cell traces from (D). Scalebars, 20 µm.

We initially characterized modErk-KTR in NIH-3T3 mouse embryonic fibroblasts to ensure our modifications had not compromised the biosensor’s Erk sensitivity ^20^. Moreover, NIH-3T3 cells divide infrequently, particular with serum starvation, and so will display minimal Cdk1-dependent effects on KTR localization ^30^. Following overnight serum starvation, cells displayed a baseline low C/N ratio indicative of low/no ERK activity (Figure 3B, C). Upon addition of 10% fetal bovine serum (FBS), we observed a rapid increase in C/N ratio for both biosensors within ~5 min, which could be inhibited within 15–30 min with MEKi (10 µM PD-0325901) (Figure 3C and Videos S2 and S3). Both displayed similar activation kinetics; however, modErk-KTR exhibited a sharper decline in C/N ratios in response to MEKi showing increased responsivity, although both displayed a similar degree of inhibition by 60 min (Figure 3B). We also observed a serum concentration-dependent response of modErk-KTR (Figure 3D, E). Addition of a high concentration of serum (10% FBS) elicited a rapid, sustained increase in C/N ratios over 1.5 hr. By comparison, a low concentration of serum (2% FBS) elicited a slower response with reduced amplitude, as well as more transient and oscillatory dynamics (Figure 3E). If no serum was added baseline C/N ratio was sustained over a similar time course with some low-level sporadic activity (Figure 3E). These data show that modErk-KTR responds to Mek/Erk activity and captures the full range of Erk signaling dynamics *in vitro*.

Next we tested the functionality of modErk-KTR *in vivo* by generating Tg(ubiP:modErk-KTR-Clover) transgenic zebrafish and firstly asking whether modErk-KTR faithfully reported the domains of Fgf/Erk signaling in DV and AP axis patterning using the binary classification of KTR localisation described above (Figure 1D). At 3.3 hpf, the reporter showed strong nuclear localization throughout the blastoderm indicating no Erk activity (Figure 4A), similarly to Erk-KTR (Figure 1D). From 3.6 hpf, unlike Erk-KTR, we observed high C/N ratios of modErk-KTR restricted to a discrete domain in marginal cells, representing the presumptive dorsal organizer (Figure 4A, Ai and Video S4). From 4.6 hpf, this domain of high C/N ratios expanded to encompass the embryonic margin, including both DCs and EVL (Figure S1A) with very few animal cells showing high C/N ratios (Figure 4B and Video S5). In addition, the width of the gradient of high C/N ratios progressively expanded from 4.6 to 5.3 hpf but remained limited to the marginal cells up until 6.0 hpf. In summary, a qualitative view of zebrafish blastulae shows that modErk-KTR recapitulates the expected patterns of Fgf/Erk activity.

**Figure 4.**
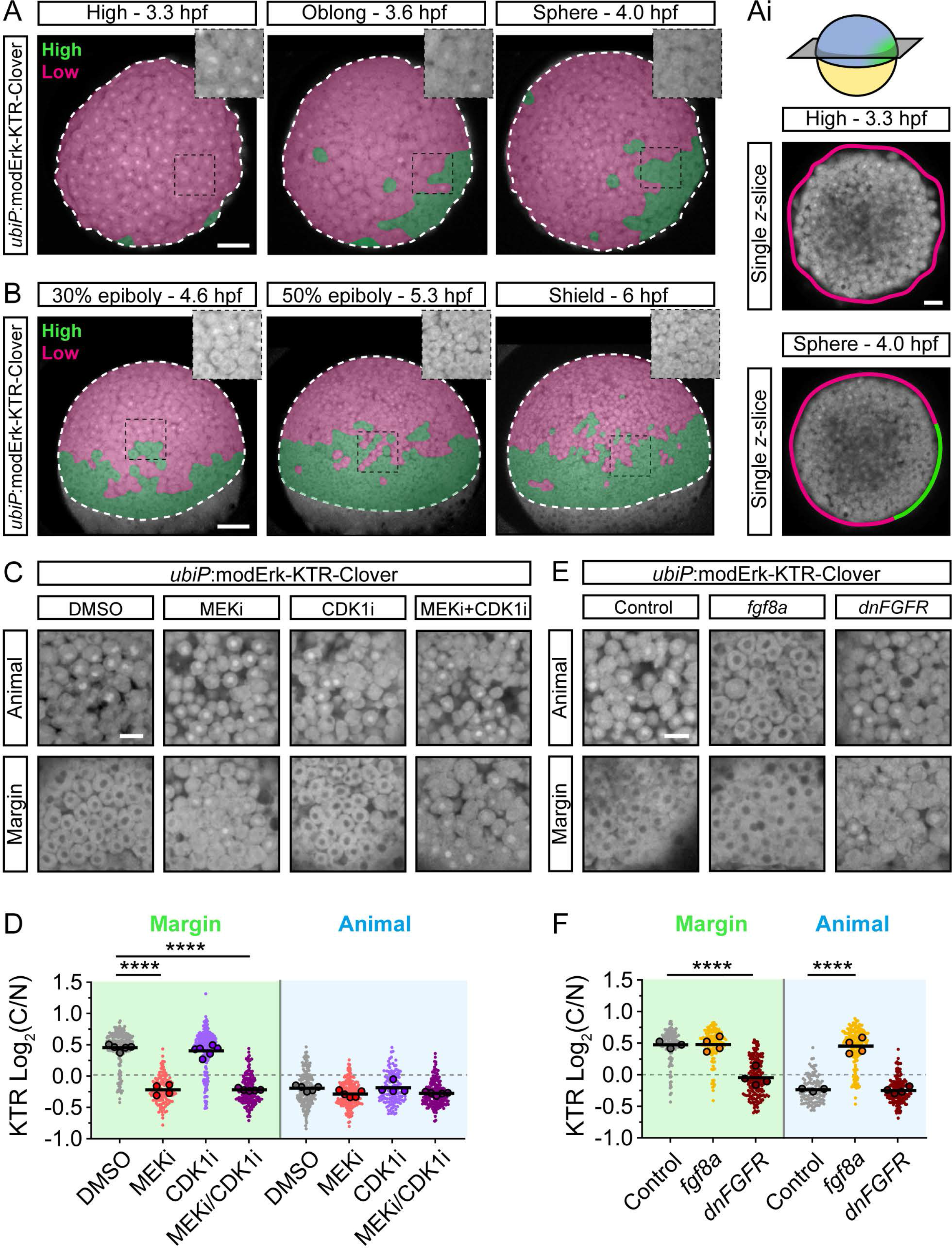
modErk-KTR specifically reports on Fgf/Erk activity in early zebrafish embryos. (A, B) Stills of live *ubiP*:Erk-KTR-Clover transgenic embryos showing a time course of reporter activity. Embryos are false colored to indicate Erk activity levels in a binary manner; green shows high and magenta shows low activity. Embryos are shown from an animal (A) or lateral (B) view. Insets show a magnified view of the region within the black box without false coloring. White dashed line, embryo proper. (Ai) Single z-slices through the centre of embryos in (A) showing Erk activity around the embryonic margin using the same color scheme as (A). (C) Live imaging of *ubiP*:Erk-KTR-Clover transgenic embryos following treatment with DMSO (control), 10 µM PD-0325901 (MEKi), 20 µM RO-3306 (CDK1i) or both MEKi and CDK1i for an hour from 4.0 hpf. (D) Quantification of Erk activity in (B) at the margin and animally. Shown are the single cell readouts of Erk activity overlayed with the per embryo averages and the overall mean for DMSO (control, n = 209 cells from 5 embryos (margin) or n = 200 cells from 5 embryos (animal)), 10 µM PD-0325901 (MEKi; n = 137 cells from 4 embryos (margin) or n = 196 from 5 embryos (animal)), 20 µM RO-3306 (CDK1i; n = 227 cells from 6 embryos (margin) or n = 133 cells from 4 embryos (animal)) or both 10 µM PD-0325901 and 20 µM RO-3306 (MEKi+CDK1i; n = 183 cells from 5 embryos (margin) or n = 194 cells from 5 embryos (animal)) treated embryos. (E) Live imaging as in (C) of embryos injected with either 25 pg fgf8a or 500 pg dnFGFR at one-cell stage. Embryos were imaged at 50% epiboly (5.3 hpf). (F) Quantification of Erk activity in (E) as in (D) for control (n = 142 cells from 3 embryos (margin) and n = 116 cells from 3 embryos (animal)), fgf8a (n = 130 cells from 4 embryos (margin) or n = 192 cells from 4 embryos (animal)) or dnFGFR (n = 168 cells from 4 embryos (margin) or n = 157 cells from 4 embryos (animal)) injected embryos. Statistical tests were one-way ANOVA with Šidák′s multiple comparisons test. Scale bars, 100 µm (A-B), 50 µm (Ai) or 20 µm (C-E); ****, p ̾ 0.0001. See also Figure S1.

To address whether the observed reporter activity is solely Mek/Erk-dependent, we monitored KTR localization at the margin and animal pole following treatment with MEKi, CDK1i or both (Figure 4C, D). Treatment from 4.0 hpf for 1 hr with MEKi caused a significant decrease in the C/N ratio at the margin in comparison with the vehicle control but had no effect on cells at the animal pole. Conversely, we observed no effect on reporter localization following addition of CDK1i, while MEKi and CDK1i together lead to a similar decrease in the C/N ratio as the MEKi alone. Importantly, we did not observe cells with high C/N ratios at the animal pole, or indeed any effect, after adding inhibitors (Figure 4C, D). This suggests that the KTR modifications have successfully extinguished the Cdk1 sensitivity in zebrafish.

To further confirm the functionality of modErk-KTR as a readout of Fgf signaling, we ubiquitously overexpressed Fgf8a or dominant negative Fgf receptor (dnFgfR) and monitored KTR localization (Figure 4E, F). Overexpression of Fgf8a resulted in no further increase in C/N ratio in the most marginal cells, but we observed a significant increase animally, with the embryos showing uniformly high C/N ratios (Figure 4F). Conversely, overexpression of dnFgfR significantly reduced C/N ratios in the most marginal cells, consistent with the reduction in P-Erk levels shown previously ^16^, while having no effect on cells at the animal pole. modErk-KTR is therefore a faithful readout of Fgf/Erk signaling in the zebrafish blastula.

We next wanted to test the utility of modErk-KTR as a general biosensor of Erk signaling in other *in vivo* contexts. Previous studies have used the Erk-KTR to monitor Erk response in muscle cell wounding at 48 hpf ^9,11^. Importantly, multinucleated muscle cells are post-mitotic and therefore, they should be free of Cdk1-dependent influence on reporter localization ^31^. We carried out a wounding assay in both Erk-KTR and modErk-KTR transgenic embryos and compared their Erk-dependent response. At homeostasis, muscle cells at 48 hpf did not display any Erk activity (Figure S1B) but rapid shuttling of the reporter out of the nucleus was observed in muscle cells surrounding the wound within 15 min of wounding with both reporters. Therefore, at this later stage of development the two reporters appear to function similarly.

Finally, we compared the two reporters in other developing tissues where there is well characterized Fgf signaling. We observed that modErk-KTR displayed clear nuclear exclusion in the developing eye and tailbud presomitic mesoderm – sites of known Fgf signaling (Figure S1C). Furthermore, we performed the same comparison in 24 hpf embryos where the midbrain-hindbrain boundary (MHB) is a source of Fgf signaling ^32^. Fgf ligands (e.g. *fgf3*) are expressed in a discrete domain at the MHB (Figure S2A) while Fgf target genes, such as *pea3*, *erm1* and *sprouty4*, are expressed in broader domains suggesting that secreted Fgf ligands act at some distance from their source ^33^. To test the functionality of the KTR reporters in this context, Erk activity was measured at 24 hpf anteriorly from the MHB in the midbrain using H2B-mScarlet-I (H2B-mSc) as a nuclear marker. Using Erk-KTR, we observed generally high C/N ratios throughout the midbrain and only observed a minimal reduction at 200 µm away from the source of Fgf ligands (Figure S2B and D). By comparison, modErk-KTR read out a steeper gradient with a stepwise feature of C/N ratios, with highest C/N ratios observed at MHB and a plateau at 75–150 µm before decreasing again at 150–200 µm (Figure S2C, D). While we were not able here to directly visualize P-Erk levels through conventional immunostaining in this tissue, we noted a similar difference (uniformly high versus graded C/N ratios) in Erk activity readout here as that in 4.0–6.0 hpf embryos (Figures 1 and 4). In conclusion, modErk-KTR displays improved Erk specificity in zebrafish embryos and can be used to monitor Erk signaling in a wide variety of developmental contexts *in vivo*.

### An improved reporter system for Drosophila embryonic and larval tissues

Recently Erk-KTR was adapted for use in Drosophila and used to monitor ERK activity in several larval and adult tissues ^34^. It was also further developed to include a histone marker (Histone 2Av (H2Av)-mCherry) produced from the same coding sequence but separated by a self-cleaving T2A peptide ^35^. The presumed equimolar concentrations of KTR:H2Av enable the readout of ERK activity by nuclear fluorescence alone in contexts where cells are densely packed and the measuring of cytoplasmic fluorescence becomes difficult ^34,35^. We asked whether modErk-KTR would offer an improvement in *Drosophila*, particularly during early development where reporter localization could be influenced by rapid cell cycles. To test this, we generated new transgenic lines with modERK-KTR and H2Av-mCherry separated by a T2A peptide (modERK-KTR-Clover-T2A-H2Av-mCherry) under the control of a *UAS* or *nanos* promoter (*nosP*) for tissue specific and maternal expression, respectively. We also generated a new transgenic line with the ERK-KTR-Clover-T2A-H2Av-mCherry under the control of *nosP* for comparison.

RTK signaling through Torso induces gradients of P-ERK at the anterior and posterior poles of the *Drosophila* blastoderm embryo, excluding the pole cells (Figure 5A) ^36^. To determine whether the KTRs enable the visualization of these signaling gradients, we imaged embryos from cell cycle (cc) 13 to cc14 (Figure 5B, C and S3A, B). We would predict if the reporters accurately read out ERK activity, we should observe nuclear exclusion at the poles and nuclear accumulation medially (Figure 5A). However, ERK-KTR shows only low-level nuclear accumulation throughout the length of the embryo with nuclear exclusion at the poles during cc13 (Figure 5B and Figure S3A). This suggests that ERK-KTR does not accurately read out ERK activity, similarly to what we observe in zebrafish embryos (Figure 1D). By cc14, coincident with significant lengthening of the cell cycle, ERK-KTR localization was consistent with the pattern of ERK activity: nuclear accumulation of ERK-KTR was evident in mediolateral regions while it was excluded from the nucleus in cells at both poles (Figure 5B and Figure S3A) ^37^. By contrast, during both cc13 and cc14 modERK-KTR displayed the predicted pattern of nuclear exclusion at both poles and nuclear accumulation mediolaterally (Figure 5C, Figure S3B and Videos S6 and S7). This suggests that modERK-KTR represents a substantially improved system for monitoring ERK signaling during *Drosophila* embryonic development.

**Figure 5.**
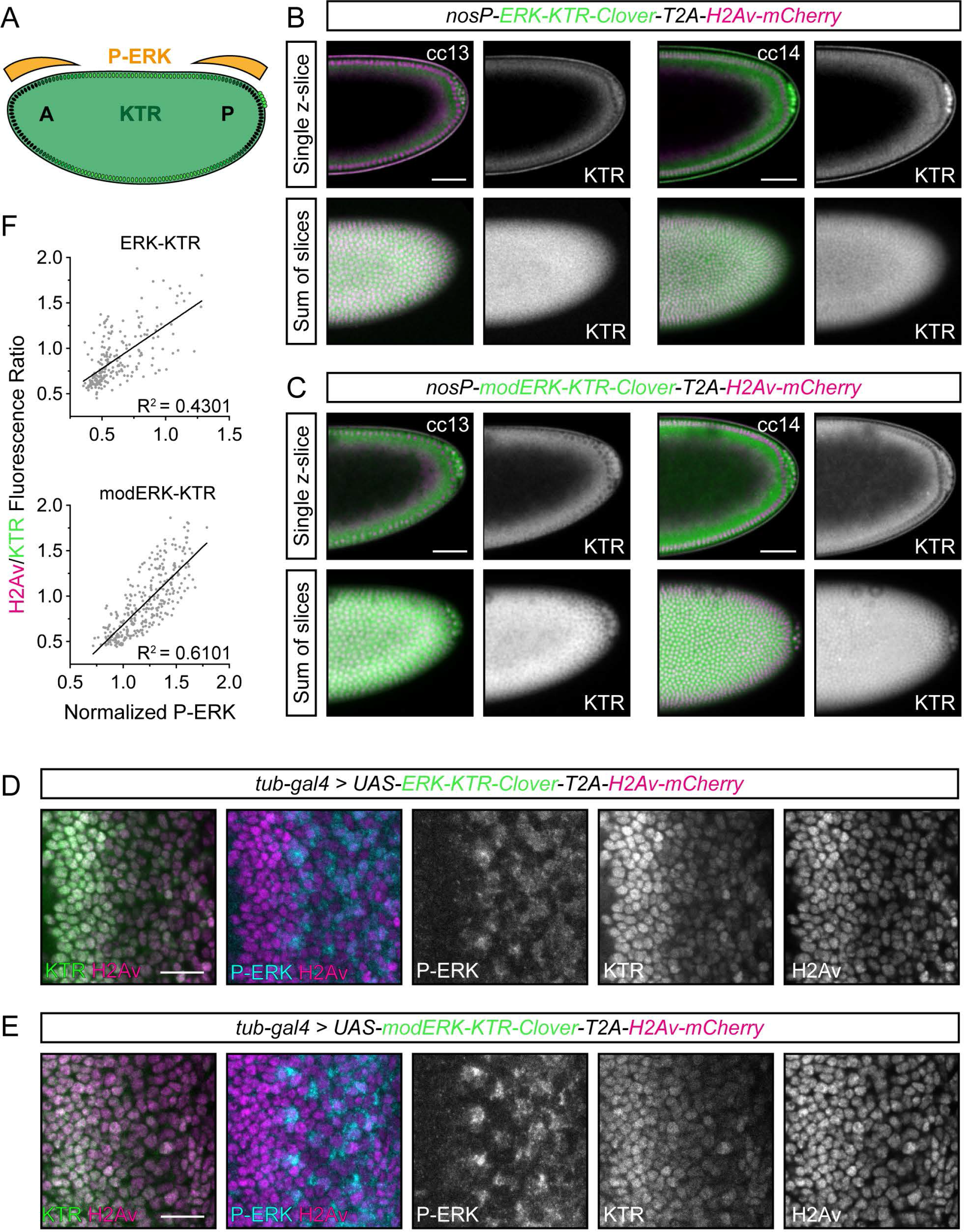
Improved Erk activity reporting with modERK-KTR in *Drosophila* embryos and larval tissue. (A) Schematic of anteroposterior Torso/ERK signaling during early *Drosophila* development where signaling is restricted to both poles of the blastoderm (shown by orange gradient), excluding the pole cells (cluster of green cells on the right hand side). Cells shown with black nuclei have high ERK signaling, whereas those with green nuclei have no ERK signaling. (B, C) Representative images of the posterior half of transgenic *Drosophila* embryos maternally expressing the original ERK-KTR (B) or modERK-KTR (C) constructs with a polycistronic H2Av-mCherry tag during cell cycles (cc) 13 and 14. Shown are both a single z-slice through the centre of the embryo (top) and a sum of slices projection of the top half of the same embryo (bottom). (D, E) Representative images of eye imaginal discs ubiquitously expressing the original ERK-KTR (D) or modERK-KTR (E) constructs with a polycistronic H2Av-mCherry tag under the control of *Tub-Gal4*. The levels of ERK activity, as read out by ERK-KTR constructs, are here compared to the levels P-ERK. (F) Quantification of (D and E) comparing levels as read out by the ERK-KTR (n = 222 cells from 3 discs) or modERK-KTR (n = 293 cells from 3 discs) constructs and P-ERK immunostaining and fitted with a simple linear regression. See also Figures S3 and S4.

We also tested whether modERK-KTR offered any improvement in third instar eye imaginal discs, where EGF–ERK activity regulates the differentiation of photoreceptors as cells pass through the morphogenetic furrow ^38^. To visualize the KTRs in eye imaginal discs, we drove ubiquitous transgene expression with *tubulin-Gal4*, performed immunostaining for P-ERK (Figure 5D, E) and compared P-ERK levels and H2Av:KTR ratios for individual cells. We found a positive correlation for ERK-KTR (R^2^ = 0.4301) as increasing levels of P-ERK correlated with a higher H2Av/KTR ratio (Figure 5F). However, the correlation between P-ERK levels and HisAv/KTR ratios for modERK-KTR showed an improved linear relationship (R^2^ = 0.6101). Indeed, we took advantage of the fact that cells in the eye disc are arrested in G1 phase in the morphogenetic furrow before some cells re-enter the cell cycle. Therefore, examining cells just posterior to the furrow and labeling S-phase cells via EdU incorporation, we could directly compare KTR localization in G1 versus S-phase cells. We focused on P-ERK-negative cells in which the KTR should be predominantly localized to the nucleus. Using ERK-KTR, we observed cells with similarly low levels of P-ERK that varied in KTR localization: EdU-positive cells (Figure S4A; white dashed lines) had lower nuclear KTR fluorescence than P-ERK-negative/EdU-negative neighbouring cells (yellow dashed lines). Thus, the cell cycle stage of cells influenced ERK-KTR localization. By contrast, modERK-KTR displayed similar nuclear enrichment in cells that were P-ERK-negative, irrespective of cell cycle phase (Figure S4A). To further examine the cell cycle dependence of ERK-KTR, we compared the readout of the reporters in the ovarian germline. In the germarium, the structure that houses the germline stem cells and their progeny, ERK signaling is restricted to the somatic cells ^39^. Early germ cells therefore provide a proliferative but completely P-ERK-negative background in which the KTRs should be localized exclusively to the nucleus. Both reporters show similar degrees of nuclear accumulation in egg chamber germ cells (Figure. S4B), but the early germ cells within the germarium show weaker nuclear enrichment of ERK-KTR compared to modERK-KTR (Figure S4C).

Together, these data show that modERK-KTR provides an improved readout of ERK activity in *Drosophila*. This highlights that cell cycle dependence of ERK-KTR localization is not a phenomenon restricted to early zebrafish development and should be considered in all proliferative cells/tissues.

### Growth of the Fgf/Erk signaling gradient in zebrafish presumptive mesendoderm

Now equipped with a specific reporter for Erk activity, we asked whether we could track the growth of the Fgf/Erk signaling gradient at the embryonic margin. Firstly, we imaged the lateral region of Tg(ubiP:modErk-KTR-Clover) embryos from 4.3 hpf, at which point Erk activity is restricted to the dorsal organizer (Figure 6A). Embryos were imaged every 5 mins for 1.5 hr and C/N ratios were measured relative to distance from the margin and presented at 20 min intervals (Figure 6B). Cells were binned into cell tiers from the margin (YSL = 0) and we noted that during this period of development, cells undergo a change in size due to proliferation (24 to 18 µm width; Figure S5A, B). Thus, the size of a single cell tier reduced over time (see methods). At 4.3 hpf, there was little to no cytoplasmic enrichment observed in the lateral region, as expected, but by 4.6 hpf the first 4 cell tiers (~100 μm) from the margin began to show slightly higher C/N ratios (Figure 6B). We note that the first cell tier (~25 μm) exhibited lower C/N ratios in comparison with cells further from margin. By 5.0 hpf, the gradient had expanded to eight cell tiers (~175 μm; Figure S5C) and by 5.3 hpf the full 10 cell tier (~200 μm; Figure S5C) gradient had formed (Figure 6B). Importantly, we observed a high degree of variability in the level of Erk activity across the gradient and at each time point (Figure S5D). This shows that the response of presumptive mesendodermal cells to Fgf signaling is heterogeneous, which is supported by our recent work showing similar heterogeneity in P-Erk levels ^19^.

**Figure 6.**
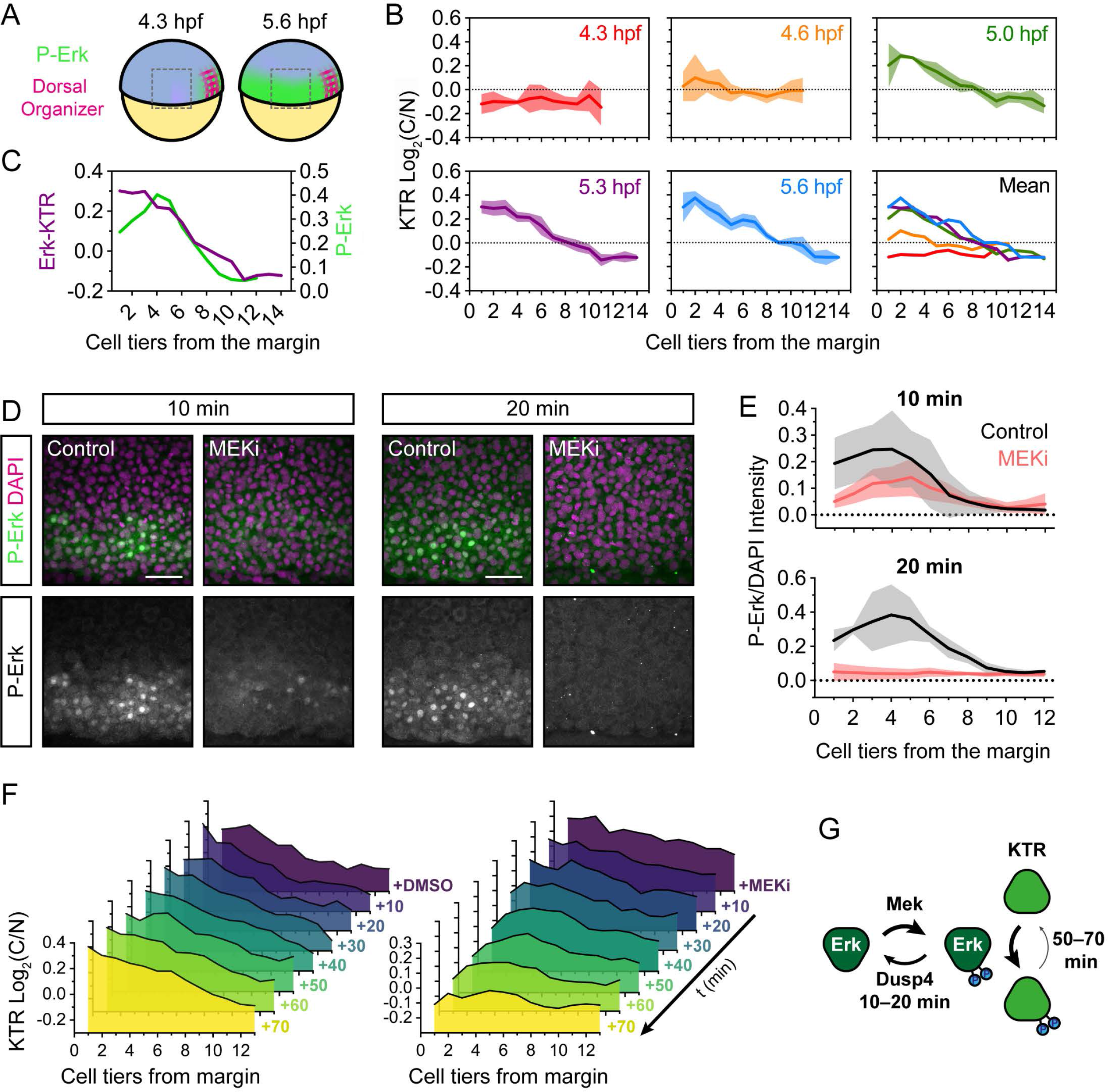
modERK-KTR reads out Fgf/Erk signaling gradient formation in real-time. (A) Schematic of the zebrafish embryo illustrating the lateral region imaged in (B) relative to the dorsal organizer (see Figure 1C). (B) Quantification of Erk activity (Log_2_(C/N)) in the lateral region of *ubiP*:modERK-KTR-Clover embryos at 20 min intervals from dome (4.3 hpf) to germ ring stage (5.6 hpf). Cells were binned based on their distance in cell tiers from the embryonic margin (0). n = 3 embryos showing the per embryo mean ± SD. Also shown is an overlay of the mean levels at each time point. (C) Overlay of the mean Erk activity (modERK-KTR) and P-Erk levels in similarly staged embryos (5.3 hpf). (D) Representative immunofluorescence images of embryos treated with DMSO (control) or MEKi (10 µM PD-0325901) for 10–20 min from 5.0 hpf before fixation. (E) Quantification of P-Erk levels from (D) in cell tiers relative to the embryo margin (0) showing mean ± SD. DMSO 10 min 5 embryos; MEKi 10 min n = 6 embryos; DMSO 20 min n = 4 embryos; MEKi 20 min n = 4 embryos. (F) Quantification of Erk activity as read out by modERK-KTR following treatment with DMSO (control) or 10 µM PD-0325901 (MEKi). Shown is the mean of n = 3 embryos per time point. (G) Schematic comparing the dephosphorylation rates of Erk and its targets. Scale bars, 50 µm. See also Figure S5.

We found that the lower levels of P-Erk in the first three cell tiers, driven by activity of the dual specificity phosphatase Dusp4, were not read out by modErk-KTR by 5.3 hpf (Figure 6C) 18. This is unlikely to be due to the sensitivity of modErk-KTR as equivalently low levels of P-Erk are read out in cell tier 6. A more likely explanation is that cell tiers 1-4 have been experiencing Erk activity for 20 min longer than cells further from the margin, therefore, modErk-KTR may read out the accumulative dose of Erk activity over time. To test this possibility, we monitored the rate of Erk deactivation (comparing P-Erk levels and modErk-KTR localization) following the addition of MEKi from 40% epiboly (5.0 hpf). This rate will be determined by the activity of both P-Erk phosphatases and phosphatases that are present that dephosphorylate the reporter.

We observed that the majority of P-Erk was dephosphorylated within 10 min of inhibitor addition and after 20 min it was completely extinguished in comparison with the DMSO control (Figure 6D, E) ^18^. Note, cell tiers 1-2 are most sensitive to MEKi and completely lose P-Erk within 10 min. Thus, the presumptive mesendoderm is therefore very sensitive to changes in Fgf/Erk signaling input (Figure 6G). Next, we monitored the rate of modErk-KTR response to MEKi. Following addition of DMSO at 40% epiboly, the gradient of KTR C/N localization built up gradually over time to an almost linear slope with highest Erk C/N ratios at the margin (Figure 6F and Figure S5E). In contrast to P-Erk, after addition of MEKi the gradient remained unchanged after 20–30 mins. In fact it was only after 40 mins that cell tiers 1–2 show a decreased C/N ratio and only after 60–70 mins that cell tiers 3–10 show a loss of Erk activity (Figure 6F and S5E). This demonstrated that the rate of modErk-KTR dephosphorylation was substantially slower than that of P-Erk in the early zebrafish embryo (Figure 6G). We note that this is slower than what we observed in NIH-3T3 cells (~30 min; Figure 3B) and therefore propose that slow KTR dephosphorylation is an inherent property of the zebrafish embryo. Indeed, this suggests that modErk-KTR reads out the cumulative dose of Erk activity in the first 1–3 cell tiers (Figure 6C). Nevertheless, the lower levels of P-Erk sensitize these cells to changes in Erk activation.

These data demonstrate the utility of modErk-KTR as a live readout of Erk activity during embryonic development and we here demonstrate the formation in live embryos of an Fgf signaling gradient *in vivo*.

### Mitotic erasure induces oscillatory Fgf/Erk signaling dynamics

Heterogeneity in Fgf/Erk signaling is apparent during zebrafish mesendodermal patterning both at the level of P-Erk ^19^ and downstream Erk activity (Figure S5D). To address how this heterogeneity arises, we injected one-cell stage embryos with *H2B-mScarlet-I* mRNA and tracked individual nuclei from ~4.6 hpf. We found that as cells approach mitosis there was a rapid (within 2–3 mins) decrease in C/N ratio (Figure 7A and C and Video S8). Post-mitosis, daughter cells initially displayed low C/N ratios, suggesting that they must reactivate signaling (Figure 7B and C). Intriguingly, we observed this same phenomenon in Drosophila at cc13 (Figure S6A). We confirmed that this is not a reporter artifact, as we also observed a loss of P-Erk in mitotic cells at the zebrafish margin (Figure 7H). This appears specific to Fgf/Erk signaling, as Nodal ligand-driven P-Smad2 is maintained throughout mitosis. These data reveal that mitotic erasure of Fgf/Erk signaling induces periods of Erk inactivity.

**Figure 7.**
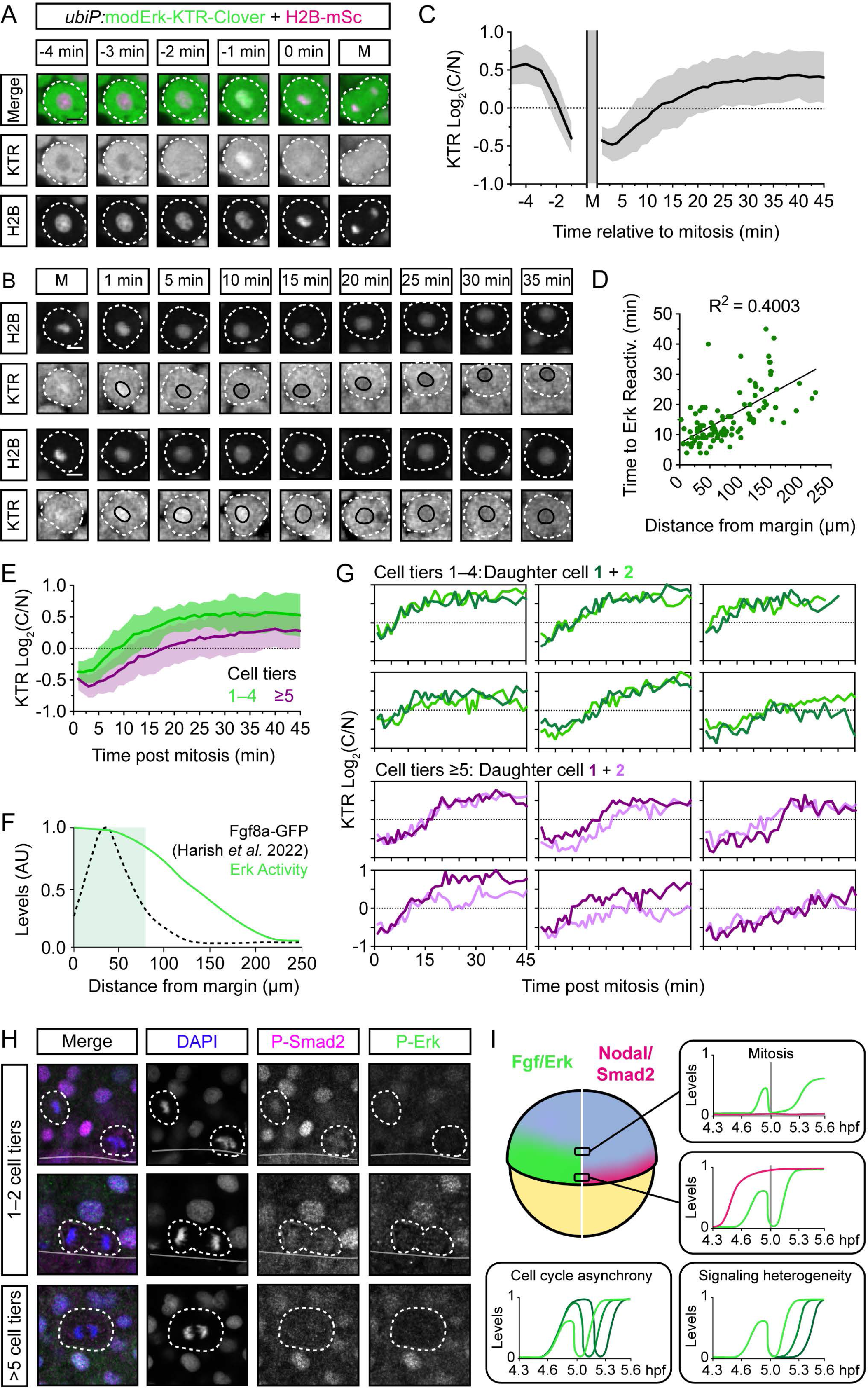
Mitotic erasure induces oscillatory Fgf/Erk signaling dynamics in the presumptive mesendoderm. (A) Representative images of a single mesendodermal cell approaching mitosis. *ubiP*:modERK-KTR-Clover embryos were injected with 25 pg His2B-mScarlet-I mRNA at the one cell stage and a lateral region of the margin was imaged from ~4.6 hpf at 1 min intervals. White dashed line labels the single cell. (B) as in (A) following two cells post-mitosis. Black line labels the nucleus. (C) Quantification of Erk activity from (A and B) following mother cells (n = 56 cells) from −5 min before mitosis and daughter cells (n = 110) +45 min after mitosis showing mean ± SD. Nuclear envelope breakdown means the KTR cannot read out Erk activity during mitosis itself. (D) Quantification of the time to Erk reactivation (log_2_(C/N) = 0.25) post-mitosis and the distance of each cell from the embryonic margin (n = 110) and fitted with a simple linear regression. (E) Comparison of Erk reactivation rates from (C) with cells binned based on the cell tier they initially start in. (F) Schematic of the Erk activity gradient, as read-out by modERK-KTR, and the extracellular levels of Fgf8a-GFP described in similarly staged embryos (~5.3 hpf) ^40^. (G) Single cell traces of sister cells post-mitosis from (E). (H) Representative immunofluorescence images of P-Erk and pSmad2 in zebrafish embryos (4.6 hpf). Mitotic cells (white dashed line) retain pSmad2 but lose P-Erk staining. (I) Model depicting how mitotic erasure of P-Erk and its target proteins induces oscillations in Fgf/Erk signaling. Both the rate of reactivation post-mitosis and the final amplitude of Erk activity are sensitive to a cell’s relative position within the Fgf signaling gradient. Coupled with cell cycle asynchrony and variability in reactivation rates, mitotic erasure also introduces heterogeneity to Fgf/Erk signaling in the presumptive mesendoderm. Scalebars, 10 µm. See also Figure S6.

There is a high degree of variability in the period of single cell oscillations around mitosis, specifically in the rate of reactivation post-mitosis (5–45 mins to Log2(C/N) >0.25) and the final amplitude of Erk activity. We next asked how these oscillatory dynamics might be influenced. We did not see a clear correlation between the Erk levels of the mother cell (−4 mins) and their daughter cells (+30 min) (Figure S6B and Video S9) which is likely due to the growth of the Fgf/Erk signaling gradient during this period. There was, however, a positive correlation between sister cells (Figure S6C–E). However, this appears to reflect the temporal Erk dynamics rather than actual levels (i.e. both sisters start with low C/N ratios and both increase over time). However, post-mitotic reactivation rates positively correlated with a cell’s distance from the margin (R^2^ = 0.4003; Figure 7D), with those cells in cell tiers 1–4 having the fastest reactivation rate and higher final levels of Erk activity (Figure 7E). The extracellular distribution of endogenous Fgf8a-GFP at the embryonic margin was recently described and ligand was shown to be concentrated around cell tiers 1–4 (ref 40) (Figure 7F and Figure S6G). Taken together with our data, this suggests that rapid post-mitotic reactivation correlates with extracellular ligand availability.

Despite the general trend toward faster reactivation rates in cell tiers 1–4 (5–20 min) versus 5–10 (10–45 mins), we still observed variability between both neighbouring cells and sister cells (Figure 7G and S6C–E) and this increased the further from the margin cells were located (Figure 7D, E and G). In addition, we noticed that cells >4 cell tiers away from the margin appear more mobile and can traverse entire cell tiers (Video S10). We therefore asked whether the final location of sister cells at +30 min post-mitosis might explain the variability in Erk activity between sister cells (Figure S6F). Indeed, there is a trend for the sister cell that moves away from the margin to exhibit lower levels of Erk activity. However, this is only apparent when sisters have separated by more than a single cell tier (>20 µm). This highlights the sensitivity of post-mitotic reactivation rate to a cell’s relative position within the Fgf signaling gradient (Figure 7E, F and Figure S6G), but also shows that there is a degree of cell autonomous heterogeneity as neighbouring sister cells can experience different reactivation rates (Figure 7G) and amplitudes of Erk activity (Figure 7G and Figure S6C–E).

In conclusion, these data show that modErk-KTR is incredibly sensitive to changes in Erk activity, such as in late G2 phase where Erk target phosphatase activity must be high. Mitotic erasure of P-Erk induces oscillations in Fgf signaling in the presumptive mesendoderm (Figure 7I) and the period and amplitude of these oscillations correlates with distance from the margin and, therefore, the source of extracellular ligands. These oscillations are a source of heterogeneity in Fgf/Erk signaling across the gradient and further noise is introduced by varying rates of post-mitotic reactivation.

## Discussion

### An improved Erk-specific biosensor

The advent of various biosensors of Erk activity has enabled the interrogation of cell autonomous and environmental factors that influence signal interpretation at the single cell level. Here we show that the original Erk-KTR also responds to Cdk1, a problem compounded by a combination of the rapid cell cycles of early development and a slow rate of Erk-KTR dephosphorylation. Despite very discrete patterns of Erk signaling, Erk-KTR localizes to the cytoplasm uniformly in most cells. By comparison, we see similar responses (Erk-KTR vs modErk-KTR) in older differentiated tissue, such as 48-hpf muscle cells, as others have reported in various contexts ^9-11,34^. This is likely due to the slower rate of proliferation in later development, and thus more infrequent peaks of Cdk1 activity, during which time Erk-KTR more faithfully reports Erk signaling. However, a direct comparison of P-Erk and KTR activity in larval Drosophila tissue shows an improvement in correlation, with similar differences in the *Drosophila* female germline and zebrafish hindbrain. Therefore, a lack of Erk specificity will likely pose a general problem when using Erk-KTR to monitor Erk signaling in proliferative cells and tissues. Indeed, a recent study identified the same problem with an Erk FRET reporter, as well as the Erk-KTR, where Cdk1-dependent reporter activity increased in late G2 phase in human colorectal cancer cells ^22^. By simply mutating the Erk docking domain to abolish Cyclin/CDK1 binding and inserting an additional Erk docking site, we successfully reduced CDK1 reporter activity to baseline, thereby substantially increasing the specificity of the reporter for Erk activity. Thus, modErk-KTR will be a valuable tool, providing an improved Erk-specific biosensor for use *in vitro* and *in vivo*.

### P-Erk vs KTR dynamics

Here we have monitored for the first time the formation of the Fgf signaling gradient in the zebrafish blastula. While we observe comparative timings to the growth of the P-Erk gradient ^18^, we note that modErk-KTR, in this context, reports on the accumulative dose of Erk activity. This results in the dampened levels of P-Erk in cell tiers 1 and 2 being saturating and driving optimal modErk-KTR nuclear exclusion by 5.3 hpf. However, we note that despite these saturating levels these cells closest to the margin are more sensitive to fluctuations in Erk signaling levels, as they exhibited a faster response to signal inhibition (MEKi) both with P-Erk levels and modErk-KTR shuttling.

We find that the rate of modErk-KTR dephosphorylation is much slower (40–70 min) than that of P-Erk (10–20 min) in the early zebrafish embryo, in contrast to more differentiated cells (~30 min) ^41^. This suggests that here there is little/no robust negative feedback down-regulating Erk target phosphorylation. We have previously shown that the P-Erk phosphatase, Dusp4, is highly expressed in the first two cell tiers from the margin and another P-Erk

phosphatase Dusp6 is also broadly expressed throughout the margin. As a result, P-Erk is rapidly lost upon inhibition of upstream activators ^18,42^. modErk-KTR dephosphorylation must be driven by different phosphatases that are present/active at lower levels during early development, but more highly expressed/active in more differentiated cells (for example, mouse embryonic fibroblasts). One likely candidate is calcineurin, a Ca^2^+-dependent phosphatase that dephosphorylates ELK1 (ref 43) and was recently shown to regulate Erk-KTR activity ^44^. This latter study also identified the ubiquitous phosphatase PP2A as a regulator of Erk-KTR localization, although PP2A targets multiple nodes in the MAPK pathway (e.g. Raf, Mek and Erk) ^45^. Indeed, the promiscuity of phosphatase catalytic subunits makes it difficult to differentiate direct versus indirect action on targets ^46^, though a broad acting phosphatase/s would be an attractive candidate for the rapid shutdown of all MAPK pathway activity.

Given the slow rate of KTR dephosphorylation in interphase cells, we propose that modErk-KTR in the early zebrafish embryo reads out the accumulative Erk signaling each cell has experienced. This is clear in the cells closest to the margin that experience dampened P-Erk levels yet display the highest KTR C/N ratios. This can be explained by their experiencing Erk signaling the longest, and the slow rate of modErk-KTR dephosphorylation. It will be important in the future to determine whether this phenomenon is shared by endogenous Erk targets and therefore provides a mechanism of ‘memory-retention’ of past Erk activity.

These data highlight a point for consideration when using KTRs: how closely linked are the phosphorylation/dephosphorylation rates of the target kinase and its KTR? In the zebrafish blastula, it is unlikely that we would be able to observe any interphase Erk dynamics due to the slow KTR dephosphorylation rate, while in other systems, such as mouse ESCs, frequent pulses (8 pulses/hr) of ERK activity can be registered using the Erk-KTR ^47^. A recent study reported a computational screen for gene circuits that could act as detectors of pulsatile Erk dynamics and identified an incoherent feedforward motif they could use to generate a synthetic reporter, called the READer circuit ^48^. Erk signaling is required to induce the circuit, but Erk must be subsequently turned-off for the expression of a fluorescent reporter. Such a circuit highlights how oscillatory signals can encode information and it would be interesting to use this system to test whether there are additional interphase Erk signaling dynamics in the presumptive mesendoderm.

## Mitotic erasure of Fgf/Erk signaling

By monitoring Erk activity at high temporal resolution (1 min intervals), we find that mitotic erasure of Erk activity and its downstream targets induces oscillations in Fgf/Erk signaling over time that are read out by modErk-KTR (Figure 7I). The consistency of Erk inactivation ~3 mins prior to mitosis suggests a link to the G2-M checkpoint and hence couples tissue growth to Fgf/Erk signaling. It has previously been noted that Erk responses are cell cycle sensitive. For example, G1-and S-phases are associated with delayed Erk activation and G2 with robust, rapid sustained signaling in yeast ^49^, while mouse ESCs display more spontaneous pulses of Erk activity early in the cell cycle ^47^. Although the mechanism of mitotic erasure is currently unknown, the Cdc25 dual-specificity phosphatases are attractive putative regulators as they are upregulated in late G2 phase to promote G2-M transition through the regulation of Cdk1 (Ref 50). While it is currently unknown whether they dephosphorylate P-Erk targets, Cdc25A has been shown to function as an ERK phosphatase in human hepatoma cells ^51,52^.

Erk signaling also plays a role in the regulation of proliferation through the control of G2/M-and G1/S-phase transitions. Inhibition of Erk activity is sufficient to arrest cells in G1 phase and slow the rate of entry into M phase ^53,54^. Conversely, hyperactivation of Erk can either enhance cell cycle entry or induce cell cycle arrest, depending on the levels of activation. Proliferation is therefore sensitive to Erk levels and mitotic erasure may play a regulatory role in Erk-dependent cell cycle progression.

Post-mitosis presumptive mesendodermal cells must re-activate Erk signaling, and we observe variability in the rate of reactivation that correlates with distance from the margin. This suggests that the rate of reactivation is sensitive to extracellular Fgf ligand availability. Indeed, it is well established that Fgf ligands elicit a concentration-dependent response, both in the amplitude and rate of Erk phosphorylation ^55,56^. There is also variability independent of distance from the margin between neighbouring and sister cells. While some of this can be attributed to the animal-marginal movement of cells in/out of the signaling domain, this is not sufficient to explain all the variability and suggests some cell autonomous heterogeneity in response to Fgf signaling. During the specification of cranial-cardiac progenitors in the invertebrate chordate, *Ciona intestinalis*, the asymmetric inheritance of internalized FGFRs enables differential daughter cell responses to uniformly distributed FGF ligand ^57,58^. This may be a common mechanism for coupling tissue growth and patterning downstream of RTK signaling ^35,59,60^ and, whether actively or stochastically driven, could lead to heterogeneity in Erk re-activation rates in the zebrafish blastula.

Like the variability in Erk activity we observe here with modErk-KTR, we have previously shown there is heterogeneity in P-Erk levels ^19^. We propose that mitotic erasure in combination with cell autonomous differences in reactivation rates, signal amplitude, and cell cycle asynchrony are all potential sources of heterogeneity and noise in Fgf/Erk signaling over time (Figure 7I). Two recent studies using the original Erk-KTR have described a role for FGF/ERK signaling dynamics in early mouse embryo patterning ^7,8^. Simon *et al.* (2020) showed that elevated ERK activity promotes primitive endoderm specification while sporadic pulses of activity are associated with epiblast specification. In addition, Pokrass *et al.* (2020) found that following mitosis FGF/ERK signaling levels diverge, which similarly dictates primitive endoderm versus epiblast differentiation through the ERK-dependent destabilisation of Nanog, a key epiblast-promoting factor. The authors did not observe mitotic erasure in these studies, but this is likely due to differences in temporal resolution. Nevertheless, these results could suggest that mitotic erasure of ERK signaling is a conserved process, that could regulate cell fate decision making. In the future, it will be interesting to investigate if/how mitotic erasure influences the interpretation Fgf signaling in the zebrafish blastula, and how this might impact embryonic patterning.

## Limitations

When using biosensors that are based on either upstream factors or substrates of a kinase of interest (KOI), it is important to establish that the biosensor and KOI activation/deactivation rates are correlated when interpreting reporter output. During early zebrafish development, the rate of P-Erk dephosphorylation is much faster than that of modErk-KTR in interphase. Importantly, this is not the case in other contexts, including around mitosis in the zebrafish blastula, when both P-Erk and the modErk-KTR are dephosphorylated within 2 min. These observations indicate that biosensors in interphase embryonic cells may be less sensitive to rapid Erk dynamics, if they are occurring, due to the stability of Erk-induced target phosphorylation.

It has also been noted previously that the KTR system alone is not ideal for use in densely packed tissues or those with non-uniformly shaped cells. To overcome this, the use of a co-expressed nuclear marker (e.g. H2Av-mCherry) has been successfully demonstrated to enable the readout of Erk activity based simply on nuclear fluorescence (Figure 5; Refs 34,35). It will therefore be useful to generate new zebrafish transgenic lines that similarly utilize this polycistronic system.

## STAR Methods

### RESOURCES AVAILABILITY

#### Lead Contact

Further information and requests for resources and reagents should be directed to and will be fulfilled by the lead contact, Caroline Hill.

#### Materials Availability

Plasmids and zebrafish lines generated in this study are maintained in the lab by the lead contact, Caroline Hill and will be made available upon request. *Drosophila* transgenic lines have been deposited in the Bloomington *Drosophila* Stock Center.

#### Data and Code Availability

This paper does not report new datasets or any original code. Any additional information required to reanalyse the data reported in this paper is available from the lead contact upon request.

### EXPERIMENTAL MODEL AND SUBJECT DETAILS

#### Zebrafish lines and maintenance

Zebrafish (*Danio rerio*) were housed in 28°C water (pH 7.5 and conductivity 500 µS) with a 15 hr on/9 hr off light cycle. All zebrafish husbandry was performed under standard conditions according to institutional (Francis Crick Institute) and national (UK) ethical and animal welfare regulations. All regulated procedures were carried out in accordance with UK Home Office regulations under project license PP6038402, which underwent full ethical review and approval by the Francis Crick Institute s Animal Ethics Committee.

#### *Drosophila* lines and maintenance

All experiments were performed in *Drosophila* melanogaster (see Key Resources Table and figure genotypes table for details of strains used). Flies were grown and maintained at 18°C and during embryo collection they were maintained at 25°C on standard *Drosophila* growth media. Embryos were collected using apple juice agar plates with additional food as above and aged to 2–4 hpf before imaging.

#### Cell culture

NIH-3T3 cells were obtained from Richard Treisman (Francis Crick Institute) and cultured in Dulbecco’s modified Eagle’s medium (DMEM) supplemented with 10% FBS and 1% Penicillin/Streptomycin (Pen/Strep). Cells have been banked by the Francis Crick Institute Cell Services, certified negative for mycoplasma and were species confirmed.

### METHOD DETAILS

#### Molecular biology and transgenesis

To generate modErk-KTR, the Erk-KTR-Clover sequence from *pDEST-ubiP:ERK-KTR-Clover-pA-Tol2* (ref 9) was codon-optimized for zebrafish with the following modifications: aga>gct (Arg>Ala) at amino acid 318 (see Figure 3A) and the addition of a C-terminal ‘tttcaattccca’ (FQFP) motif. The modified sequence was subcloned into the BamHI sites of *pDEST-ubiP:ERK-KTR-Clover-pA-Tol2* to generate *pDEST-ubiP:modErk-KTR-Clover-pA-Tol2*. Transgenic zebrafish were generated by injecting the plasmid into zebrafish embryos, which was randomly inserted into the genome using Tol2 recombinase-mediated transgenesis. To generate *pCS2-mScarletI-H2B*, mScarlet-I was amplified from *pmScarlet-i_C1* (Addgene, # 85044) and *H2B* was amplified from *pCS2-mKeima-H2B* (a gift from Nancy Papalopulu) with an N-terminal GS-linker and inserted into *pCS2* at *EcoRI* and StuI sites.

The modERK-KTR sequence with T2A-His2Av-mCherry as previously used 34 was codon-optimized for *Drosophila* and synthesized by Thermo Fisher Scientific GeneArt, then subcloned using EcoRI and SalI into the EcoRI and XhoI sites of *pUASt-attB* (ref 34). The insert was excised from pUASt-attB-modERK-KTR-T2A-His2Av-mCherry and subcloned into the NotI and NheI sites of pCasper-nosP-HA-brat-attB, replacing HA-brat (a gift from Hilary Ashe). This generated *pUASt-modERK-KTR-T2A-H2Av-mCherry-attB*, *pNosP-ERK-KTR-T2A-H2Av-mCherry-attB* and *pNosP-modERK-KTR-T2A-H2Av-mCherry-attB*. Transgenic flies were generated by injection of the plasmids into fly embryos carrying an attP2 landing site and integrated using ΦC31 integrase. Injections were carried out by BestGene Inc or the Crick Fly Facility. All transgenic lines have been deposited and are available from the Bloomington *Drosophila* Stock Center, Bloomington, Indiana, USA.

#### mRNA injection of zebrafish embryos

Capped RNA for injection was transcribed using the mMessage mMachine Sp6 or T7 kit (ThermoFisher Scientific) followed by LiCl precipitation. For live imaging experiments, zebrafish embryos were injected with 25 pg *H2B-mScarlet-I* mRNA at the one-cell stage. For overexpression experiments, embryos were injected with 50 pg *fgf8a* or 500 pg *dnFGFR* mRNA.

#### Cell culture

NIH-3T3 cells were grown in a glass bottom 35 mm MaTek dish and transfected with the *pDEST-ubiP:ERK-KTR-Clover-pA-Tol2* or *pDEST-ubiP:modERK-KTR-Clover-pA-Tol2* using FuGene (Promega) according to the manufacturer’s instructions. Prior to imaging, cells were incubated in DMEM with 0.5% FBS overnight to ensure baseline ERK activity. Erk activity was induced by addition of 10% FBS.

#### Live imaging

Zebrafish embryos were collected and maintained at 28°C until 3.5 hpf. Embryos were mounted in their chorion in 1% low melting agar (Sigma) on a glass bottom 35-mm MaTek dish and bathed in embryo media with or without chemical inhibitors (see below). Embryos were oriented manually to ensure a lateral view and to exclude the dorsal region, which experiences early Fgf/Erk activity at 4 hpf. Embryos were imaged on a Leica SP8 inverted confocal microscope using an HC PL APO CS2 20x/ 0.75 IMM objective at 28 °C with the following confocal settings, pinhole 1 airy unit, scan speed 400 Hz unidirectional, format 512 × 512 pixels at 8 bit. Images were collected using hybrid detectors and an argon and 561 nm lasers with 2x line averaging and z-slices taken at 2 µm intervals every 1 min (Figure 7) or 5 min (Figure 6). Imaging of the dorsal hindbrain at 24 hpf was carried out as described previously ^61^.

Live imaging of *Drosophila* embryos was carried out as described 62. Embryos were dechorionated in bleach and positioned laterally on top of a coverslip (No. 1, 18 × 18 mm) thinly coated with heptane glue. A drop of halocarbon oil mix (4:1, halocarbon oil 700: halocarbon oil 27)) was placed in the middle of a Lumox imaging dish and two coverslips (Nr. 0, 18 × 18 mm) were placed on either side of the oil drop. The coverslip with the embryos attached was then inverted into the oil, sandwiching the embryos between the imaging dish membrane and the coverslip. Embryos were imaged on a Leica SP8 inverted confocal microscope using an HC PL APO CS2 20x/ 0.75 dry objective at 25°C with the following confocal settings, pinhole 1 airy unit, scan speed 400 Hz unidirectional, format 512 × 512 pixels at 8-bit. Images were collected using hybrid detectors and an argon and 561 nm lasers with 1x line averaging and z-slices taken at 2 µm intervals every 3 min.

Live imaging of NIH-3T3 cells was performed as described above on a Leica SP8 inverted confocal microscope using an HC PL APO CS2 20x/ 0.75 IMM objective at 37°C and 10% CO_2_. Images were collected with 2x line averaging and z-slices taken at 1 µm intervals every 1.5 min.

#### Zebrafish wounding

Embryos at 48 hpf were immobilized with tricaine (0.08 mg/ml) in E2 buffer and mounted laterally in 1% low melting agar on a glass bottom 35-mm MaTek dish. Wounding was achieved by manually puncturing the muscle with a glass needle.

#### Fluorescence in situ hybridization (FISH) and immunofluorescence (IF)

Combined FISH and IF was performed with the RNAscope^®^ 2.0 Assay using the Multiplex Fluorescent Assay v2 (ACDBio) as previously described ^63^ with minor modifications. Briefly, after fixation in 4% paraformaldehyde (PFA), followed by incubation overnight in methanol, embryos were rehydrated and incubated with Dr-gsc (427301-C3, ACDBio), Dr-*fgf8a* (559351-C2, ACDBio) and/or Dr-*fgf3* (850161-C4, ACDBio) probes at 40°C overnight. Embryos were then washed in 0.2x saline sodium citrate/0.01% Tween 20 (SSCT) and re-fixed in 4% PFA for 10 min followed by washes with SSCT. First, they were incubated with two drops of the Amp1 and Amp2 solution at 40°C for 30 min and then incubated with two drops of Amp3 at 40°C for 15 min. After an additional washing step, embryos were incubated with two drops of the Multiplex FL V2 HRP-C2, -C3 or -C4 at 40°C for 15 min. After a last series of washes in SSCT, embryos were washed in PBS/ 0.1% Tween-20 (PTW) and processed for the staining. Like conventional FISH, embryos were incubated with tyramide (Sigma) coupled with fluorescein-NHS ester (Thermo Scientific, #46410), Cy3 mono NHS ester (Sigma, #PA13101) or Cy5 mono NHS ester (Sigma, #PA15101) in PTW in the dark. To allow HRP detection, 0.001% H_2_O_2_ was added to the reaction and embryos were incubated for 30 min, also in the dark. The embryos were then extensively washed in PBS/1% Triton X-100 (PBTr) and incubated in acetone at −20°C. After that, embryos were incubated for 2 hr in PBTr with 10% FBS before incubation with antibodies against P-Erk overnight at 4°C. Antibody binding was detected with HRP-conjugated anti-mouse secondary antibodies and signal was developed as above for RNA detection.

IF for P-Smad2 and P-Erk was performed as described ^18^ with minor modifications. Embryos were rehydrated into PBTr before incubating in acetone at −20°C. Embryos were blocked in 1% PBTr and 10% FBS, before incubating with antibodies against pSmad2 (Cell Signaling Technology, # 8828, 1:500) and/or P-Erk (Sigma, M8159, 1:500) at 4°C overnight. Antibody binding was detected with Anti-Rabbit Alexa Fluor 488 (ThermoFisher Scientific, # A-21206, 1:1000) and Anti-Mouse Alexa Fluor 594 (ThermoFisher Scientific, # A-21203, 1:1000).

In all cases, zebrafish embryos were extensively washed and DAPI was used at 1:1000 in PTW for 15 min at room temperature. Embryos were then mounted in 1% low melting agarose on a glass bottom 35-mm MaTek dish and manually oriented.

For the IF of *Drosophila* eye imaginal discs, wandering 3^rd^ instar larvae were dissected in Schneider’s insect medium (ThermoFisher, # 21720-024) and incubated in 10 µM EdU (5-ethynyl-2’-deoxyuridine) in Schneider’s medium while shaking for 30 min. After incubation, samples were fixed for 15 min in 4% paraformaldehyde in 10 mM Tris-HCl (pH 6.8), 180 mM KCl, 50 mM NaF, 10 mM NaVO4, and 10 mM β-glycerophosphate, then washed twice in 0.5% PBTr for 30 min. Samples were blocked in 0.2% PBTr and 1% FBS for 1 hour, then incubated overnight at 4°C in rabbit anti-phospho-ERK antibody (Cell Signaling Technology, # 9101, 1:200). The samples were then washed twice for 30 min in 0.5% PBTr and 1% FBS and subsequently incubated in secondary antibody for 2 hr at room temperature, before being washed in 0.2% PBST for 30 min, then incubated for 30 minutes in 2.5 µM AZ dye 405 picolyl azide (Click Chemistry Tools), 0.1 mM THPTA, 2 mM sodium ascorbate, and 1 mM CuSO4. Finally, the samples were washed twice in 0.2% PBTr for 15 minutes and mounted on microscope slides with Vectashield medium (H-1000, Vector labs).

For imaging *Drosophila* ovaries, adult females were raised with males for 3–7 days post-eclosion prior to dissection in order to promote normal reproductive health. Ovaries were dissected in Schneiders insect medium and fixed in 4% PFA for 15 mins. They were extensively washed in 0.1% PBTr and DAPI was used at 1:1000 for 15 min at room temperature. Ovaries were mounted on a microscope slide in Prolong Gold Antifade.

All FISH and IF samples were imaged on a Leica SP8 inverted confocal microscope using either a HC PL APO CS2 20x/0.75 DRY objective or 10x DRY objective. Imaginal discs were imaged on a Zeiss LSM880 with a 40X objective.

#### Pharmacological inhibitors

For drug treatments, the inhibitors PD-0325901 and RO-3306 were dissolved in DMSO and directly diluted in embryo or cell culture medium at 10 μM (PD-0325901) and 20 μM (RO-3306) respectively. Embryos were maintained at 28°C and the time of treatment and durations are specified in the Figure legends.

### QUANTIFICATION AND STATISTICAL ANALYSIS

#### Image analysis

To quantify single cell Erk activity (Figure 2–4), a 5–6 pixel width region of interest was drawn in the centre and periphery in Fiji to measure the nuclear and cytoplasmic mean intensities, as illustrated in Figure 2A. These were used to calculate the log2(cytoplasmic/nuclear) to give a linear readout of Erk activity. To track cells pre- and post-mitosis, H2B-mScarletI nuclear signal was used to track single cells manually and XY coordinates were also measured relative to the margin (Y = 0 μm).

To quantify Erk activity across the entire Fgf signaling gradient, a lateral view of the embryo was oriented relative to the margin (Y = 0 μm) and region of interest is drawn to exclude the EVL. H2B-mScarletI was used to generate a nuclear mask with unique identifiers using CLIJ ^64^. This was performed on 4–5 single z-slices at 15–20 μm intervals to capture up to 300 μm from the embryonic margin whilst ensuring no overlap between slices. The nuclear mask was dilated by 2 pixels and the original nuclear mask subtracted to generate a cytoplasmic mask with the same ID. The nuclear and cytoplasmic masks were then used to measure mean intensity and XY coordinates. Cells were then grouped into either 20 or 25 μm bins, depending on the stage of development, to determine the number of cell tiers away from the margin. In the brain, nuclei were too closely clustered and therefore Erk activity was manually measured as above.

To quantify P-Erk levels, DAPI was used to generate a nuclear mask and measure P-Erk and DAPI intensity as well as XY coordinates in Fiji. P-Erk levels are presented relative to DAPI intensity and presented relative to the margin as above.

Cell width was measured by manually drawing a line across the centre of cells in the animal-margin axis using the line drawing tool in Fiji.

Analysis of *Drosophila* imaginal discs was carried out in Icy (v2.4.2.0). For each cell, the focal plane containing the largest nuclear diameter was identified. Using the freehand ROI tool, an approximate outline of the nucleus was drawn using the His2Av-mCherry signal. In areas where nuclei were closely packed, adjacent focal planes were used to determine the most suitable ROIs while avoiding overlapping pixels with adjacent cells. The mean intensities of each channel were then measured for each cell and used to obtain the mCherry:Clover ratio as a readout of KTR activity.

#### Statistical Analysis

Statistical comparisons were performed using two-tailed Student’s t-tests, one-way ANOVA with multiple comparisons or paired t-test as indicated in the figure legends using GraphPad Prism and Microsoft Excel. Statistical significance was assumed by p < 0.05. Individual p values are indicated, and data are represented by the mean and standard deviation unless otherwise specified. A linear regression in JMP was used for statistical analysis and fitting a line to the imaginal disc data.

## Acknowledgments

We are very grateful for the support for this work from the Francis Crick Institute Aquarium, the Fly Facility, and the Light Microscopy Facility. We thank Andrew Sharrocks for advice on KTR modifications, Nic Tapon for collaborative introductions, Hilary Ashe, Martin Distel and Nancy Papalopulu for reagents, Joachim Kurth for fly injections and Birgit Aerne for help with cloning. We thank Toby Andrews, Rashmi Priya, MC Ramel, Nic Tapon and all the members of the Hill lab for helpful discussions and very useful comments on the manuscript. This research was funded in part, by the Wellcome Trust [FC001095]. For the purpose of Open Access, the corresponding author has applied a CC BY public copyright licence to any Author Accepted Manuscript version arising from this submission. This work was supported by the Francis Crick Institute which receives its core funding from Cancer Research UK (FC001095), the UK Medical Research Council (FC001095), and the Wellcome Trust (FC001095), and a UCL/Wellcome Trust Institutional Support Fund Grant (204841/Z/16/Z) and an MRC Career Development Award (MR/P009646/2) to MA.

## Author contributions

S.G.W. and C.S.H. conceived and designed the study. S.G.W. planned, performed the experiments and analyzed the data with help from L.G. and P.A.R. C.S.H and M.A. provided supervision and funding for the study. S.G.W. and C.S.H. wrote the manuscript.

---

**Figure S1.**
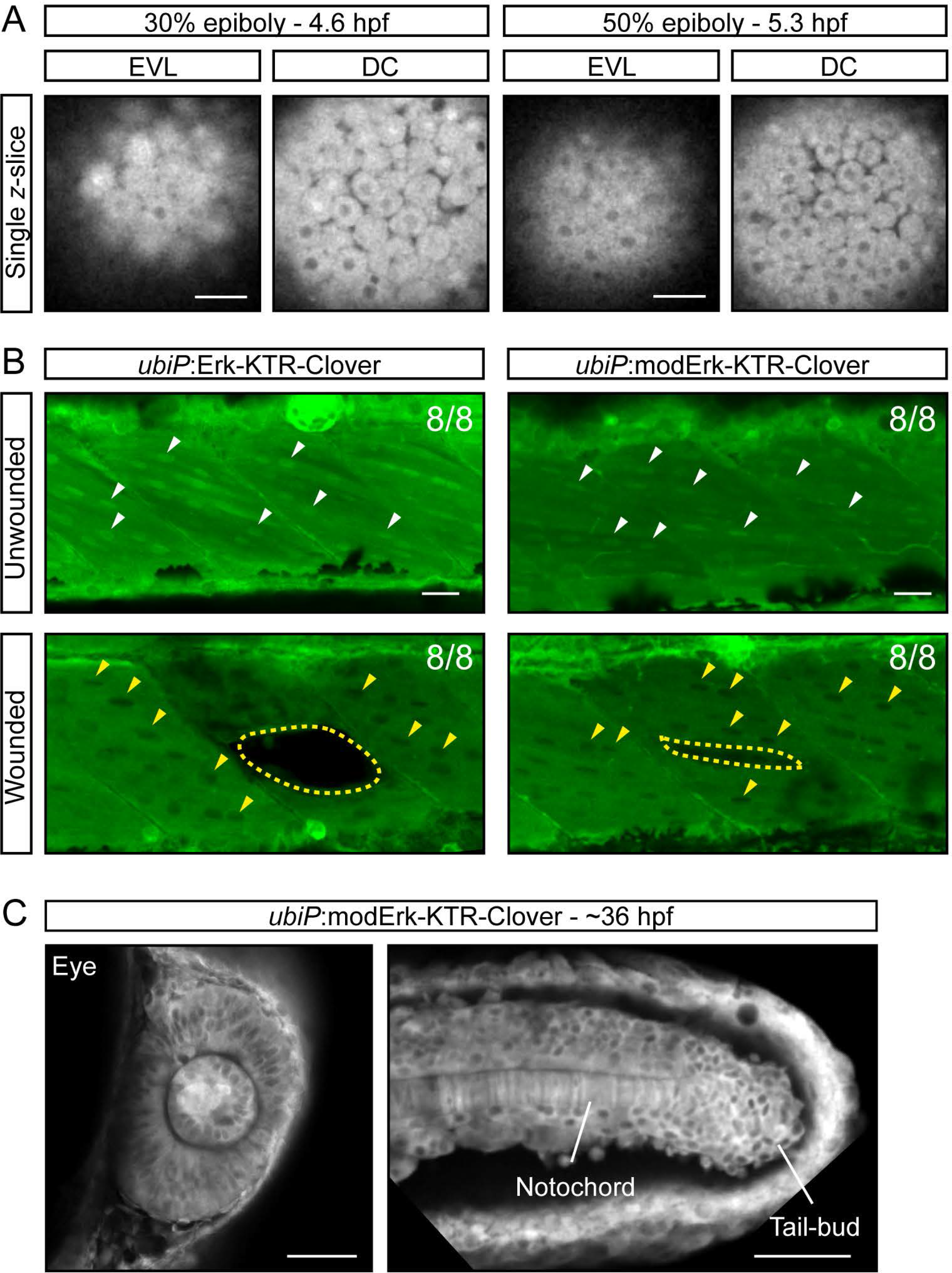
Wounding-induced Erk signaling is readout by Erk-KTR and modErk-KTR. (A) Single z-slices showing Erk-KTR activity in the EVL and DCs from the indicated embryos shown in Figure 4B. (B) Representative images of 48-hpf *ubiP*:Erk-KTR-Clover and *ubiP*:modErk-KTR-Clover zebrafish embryos with and without muscle wounding. Muscle cells typically display no Erk activity (white arrowheads). However, upon wounding the muscle the surrounding cells rapidly (~15 min) display high Erk activity (yellow arrowheads). Dashed line labels site of wound. (C) Representative images of *ubiP*:modErk-KTR-Clover zebrafish embryos in other contexts where Fgf/Erk signaling is well characterized, including the developing eye and presomitic mesoderm. Erk activity is not observed in the notochord. Scalebars, 20 µm (A, B) or 50 µm (C). Related to Figure 4.

**Figure S2.**
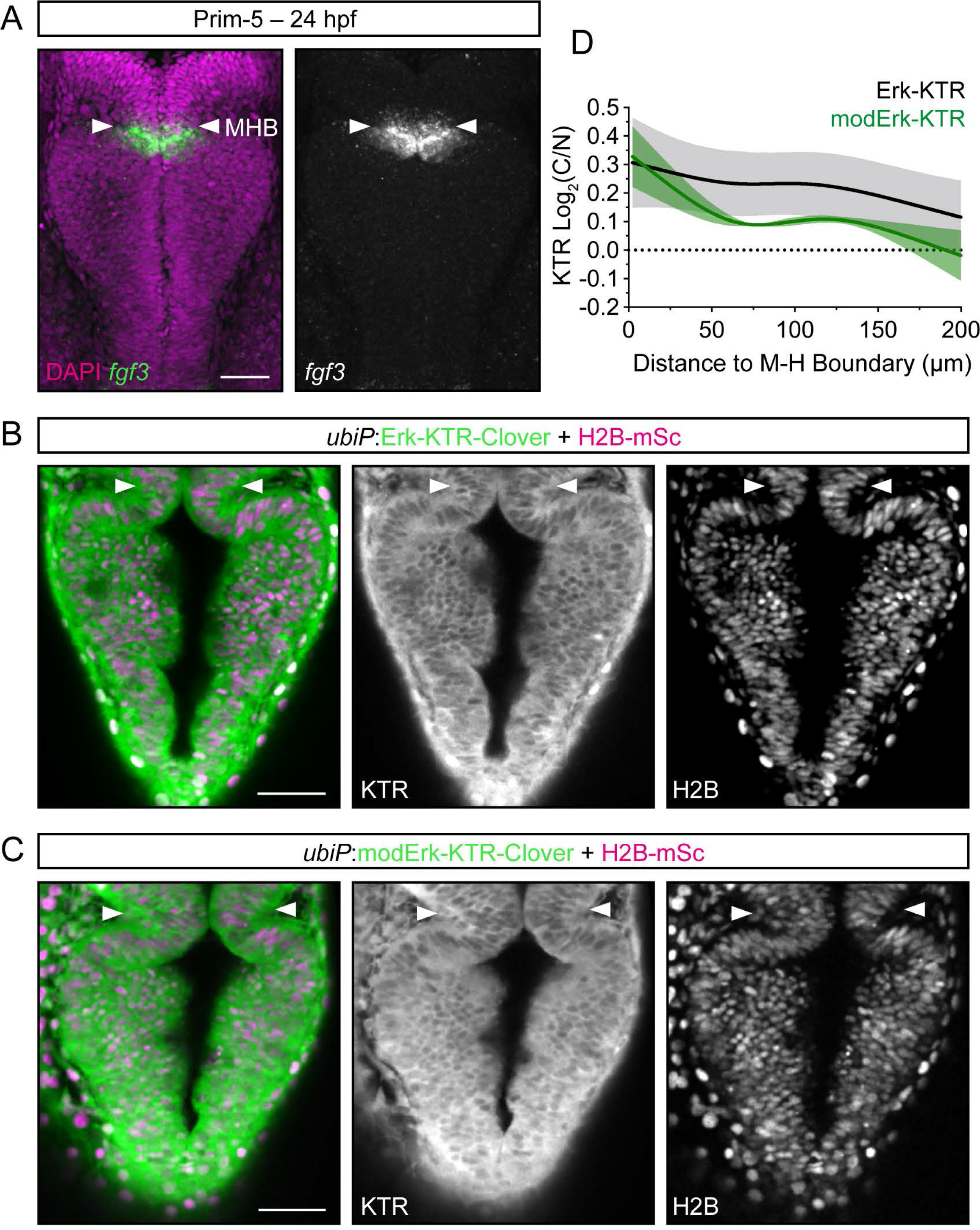
Visualization of an Fgf/Erk signaling gradient at the zebrafish midbrain-hindbrain boundary. (A) RNAscope showing expression of fgf3 at the midbrain-hindbrain boundary (MHB; white arrows) at 24 hpf. (B, C) Representative images showing a single plane through the dorsal midbrain of a *ubiP*:Erk-KTR-Clover (C) and a *ubiP*:modErk-KTR-Clover (D) zebrafish embryo. White arrows, MHB. (D) Quantification of the levels of Erk activity relative to distance from the MHB (n = 3 embryos each; mean ± SD). Scalebars, 50 µm.

**Figure S3.**
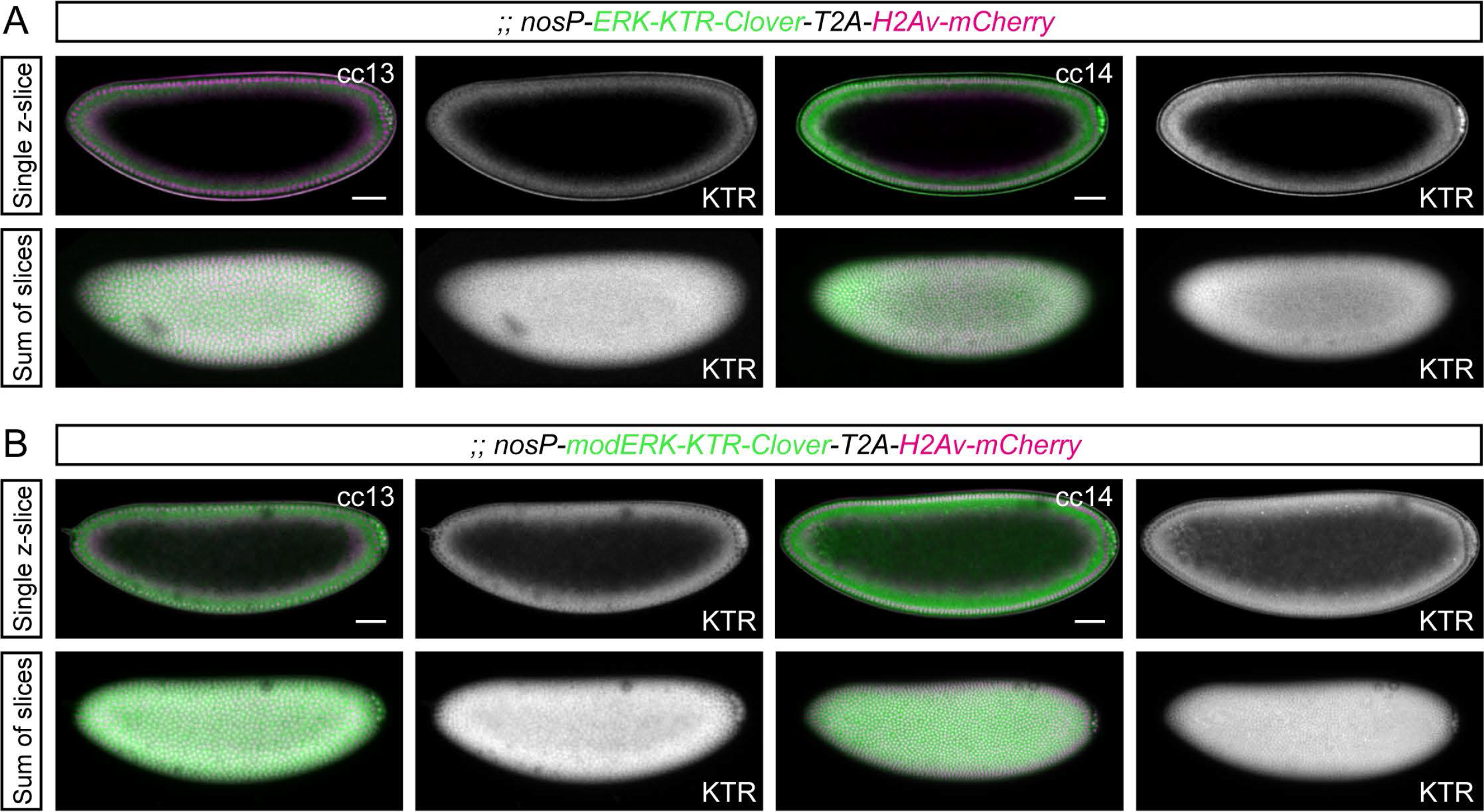
Comparison of anteroposterior Torso/Erk signaling readout in *Drosophila* embryos. (A and B) Whole embryo views of the images in Figure 5A and B showing the readout of Erk activity by the original ERK-KTR (A) and modERK-KTR (B) reporters during cell cycles (cc) 13 and 14. Shown are both (top) a single z-slice through the centre of the embryo and (bottom) a sum of slices projection of the top half of the same embryo. Related to Figure 5.

**Figure S4.**
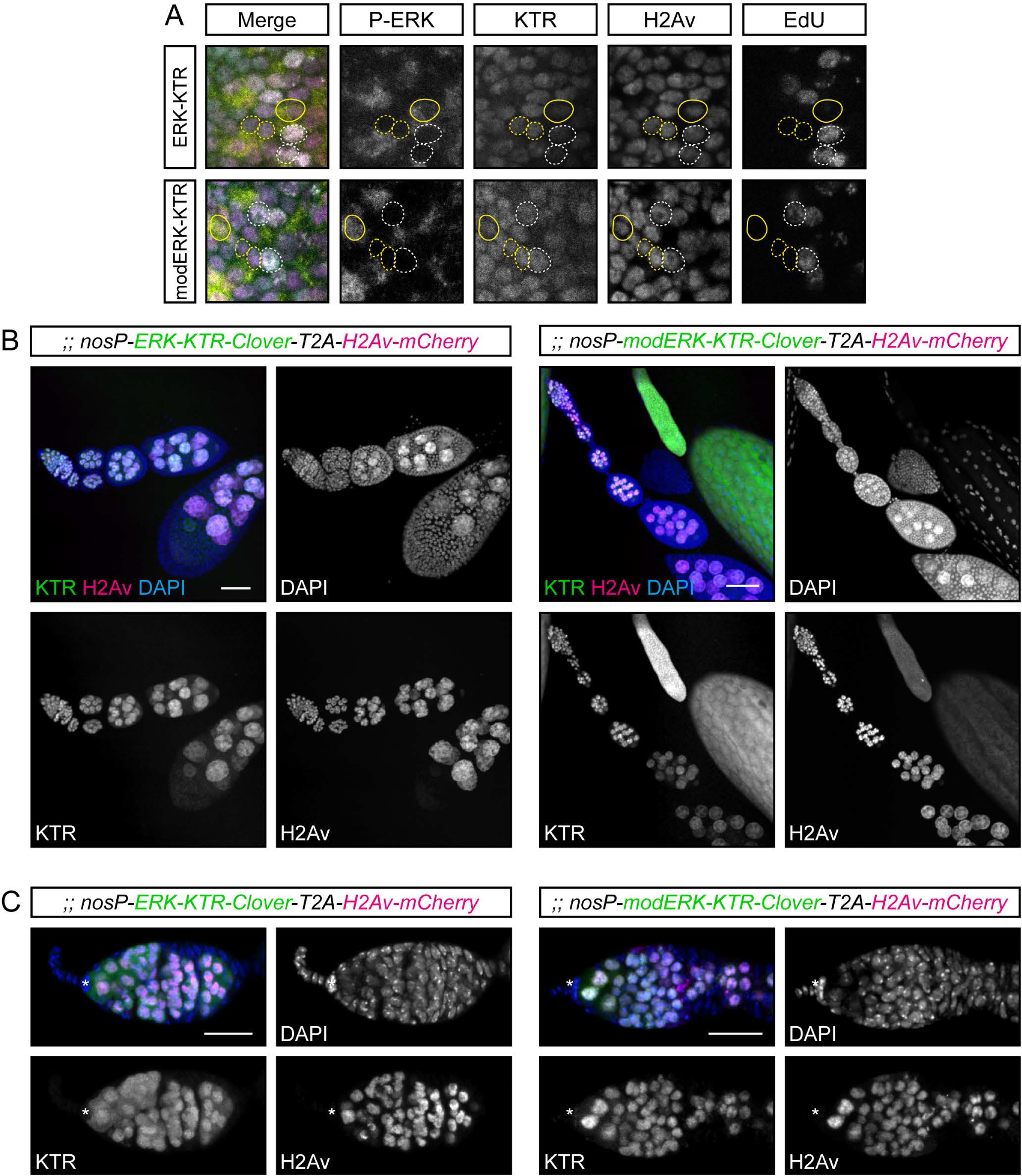
Comparison of the KTR constructs in the ovarian germline. (A) Immunofluorescence images of P-ERK levels in both KTR constructs in eye imaginal discs. Dashed lines mark P-ERK negative cells that are either EdU positive (white) or negative (yellow). Yellow lines mark P-ERK positive cells. (B) Representative images of fixed transgenic *Drosophila* ovarioles stained with DAPI to visualize the somatic tissue. nosP drives expression throughout the ovarian germline. (C) Representative images of fixed germaria as in (B). Scalebars, 50 µm (B) or 20 µm (C). Related to Figure 5.

**Figure S5.**
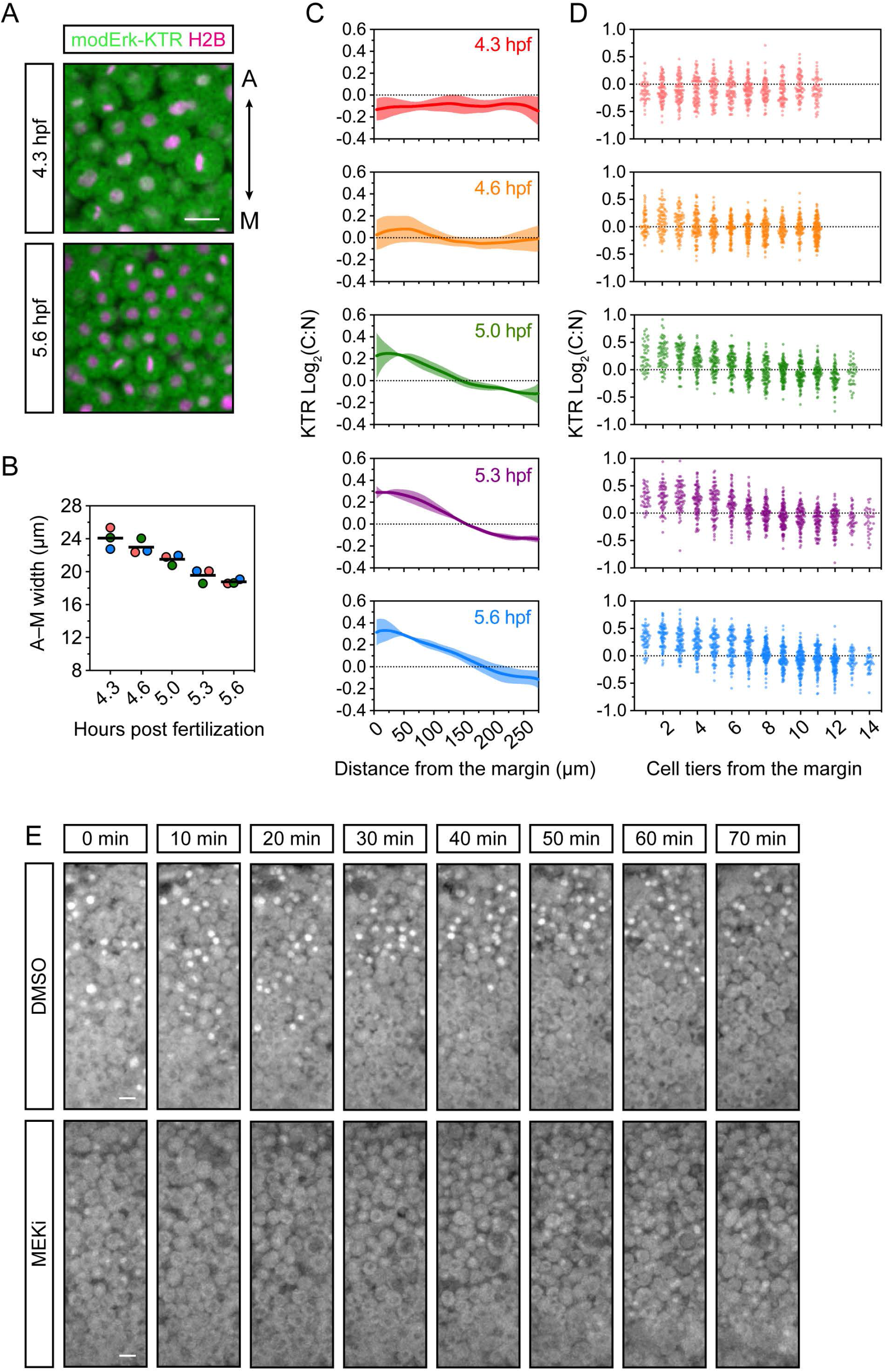
Interpreting the Fgf/Erk signaling gradient and heterogeneity in signaling levels. (A) Representative images of a single *ubiP*:modErk-KTR-Clover zebrafish embryo injected with 25 pg H2B-mScarlet-I mRNA at the one-cell stage and imaged at the indicated times of development. Embryos were imaged laterally and oriented for measuring cell width in the animal-margin (A-M) plane. (B) Quantification of A–M width from (A) at the indicate times during development. n = 40 cells per embryo, n = 3 embryos per timepoint, black line shows the mean. (C) Quantification of Erk activity (log_2_(C/N)) in the lateral region of *ubiP*:modERK-KTR-Clover embryos (Ai) at 20 min intervals from dome (4.3 hpf) to germ ring stage (5.6 hpf), relating to Figure 6B. Erk activity is shown relative to distance from the embryonic margin, (D) Quantification of Erk activity as in Figure 6B, showing all data points. (E) Representative images of lateral views of *ubiP*:modErk-KTR-Clover zebrafish embryos imaged at 5-min intervals following addition of control (DMSO) or 10 µM PD-0325901 (MEKi). Scalebars, 20 µm. Related to Figure 6.

**Figure S6.**
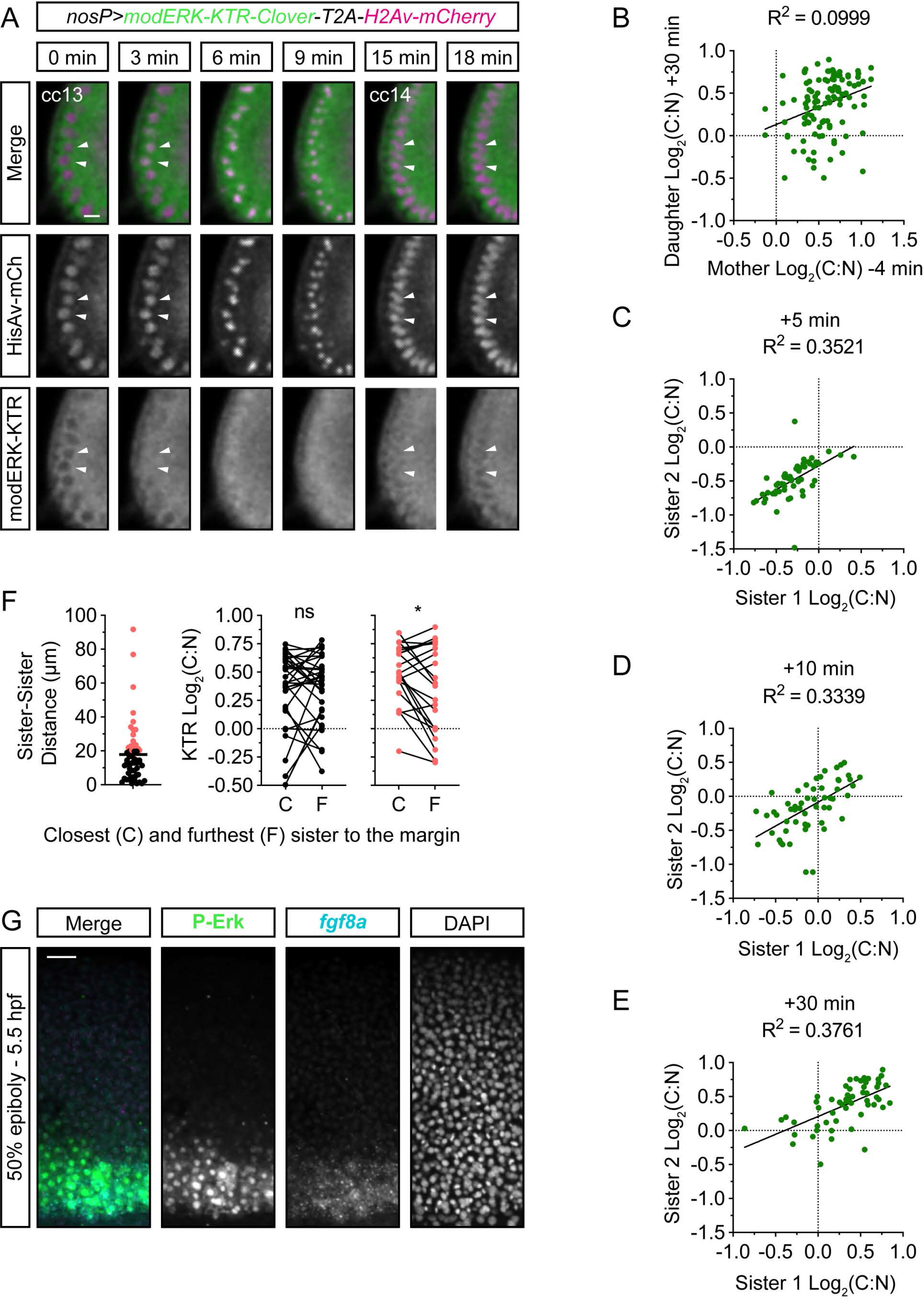
Heterogeneity in sister cell Erk activity. (A) Time course showing the anterior pole of a nosP>modERK-KTR-Clover-T2A-H2Av-mCherry *Drosophila* embryo. The embryo was imaged every 3 min and Erk activity monitored around mitosis. White arrowheads label the first two cells to undergo mitosis. (B) Quantification of the levels of Erk activity in the mother cell (n = 55) at −4 min pre-mitosis and daughter cell (n = 110) at +30 min post-mitosis and fitted with a simple linear regression. (C–E) Quantification of sister cell Erk activity levels at +5 min (C), +10 min (D) and +30 min (E) post-mitosis and fitted with a simple linear regression. (F) Quantification of the final distance between sister +30 min post-mitosis in the animal-marginal plane. (Left) Sisters that have drifted at least a single cell tier away (≥ 20 µm) were compared to those that remain in proximity (black). (Right) Pair-wise comparison of Erk activity between the sister closest to the margin (C) and the sister furthest from the margin (F). Paired t test; ns, non-significant (p = 0.5501); *, p = 0.0173. (G) Combined immunofluorescence and RNAscope showing a P-Erk and fgf8a expression at the zebrafish embryonic margin at 5.5 hpf. These are zooms of the images shown in Figure 1C. Scalebars, 10 µm (A) or 50 µm (G). Related to Figure 7

**Video S1. Erk-KTR displays off-target homogeneous activity.** Maximum projection of an animal-lateral view of a developing Tg(*ubiP*:Erk-KTR-Clover) zebrafish embryo from sphere stage (4.0 hpf) for 182 min.

**Video S2. Erk-KTR activity in response to serum in NIH-3T3 cells.** A representative example of an NIH-3T3 cell following overnight serum starvation and addition of 10% FBS at t = 0 min and addition of 10 µM MEKi at t = 30 min.

**Video S3. modErk-KTR activity in response to serum in NIH-3T3 cells.** A representative example of an NIH-3T3 cell following overnight serum starvation and addition of 10% FBS at t = 0 min and addition of 10 µM MEKi at t = 30 min.

**Video S4. modErk-KTR activity in an early zebrafish blastula.** Maximum projection of a animal view of a developing Tg(*ubiP*:modErk-KTR-Clover) zebrafish embryo from high stage (3.3 hpf) for 80 min.

**Video S5. modErk-KTR activity in an epiboly stage zebrafish embryo.** Maximum projection of an animal-lateral view of a developing Tg(*ubiP*:modErk-KTR-Clover) zebrafish embryo from 30% epiboly (4.6 hpf) for 128.8 min.

**Video S6. Anteroposterior Erk signalling in *Drosophila*.** Sum of slices projection of the lateral view of a cc12 stage nosP-modERK-KTR-T2A-His2Av-mCherry transgenic *Drosophila* embryo imaged every 3 min through to cc14.

**Video S7. Anteroposterior Erk signalling in *Drosophila*.** A single z-slice through the centre of the embryo in Video S7 showing a nosP-modERK-KTR-T2A-His2Av-mCherry transgenic *Drosophila* embryo imaged every 3 min through to cc14.

**Video S8. Mitotic erasure of Erk activity in zebrafish presumptive mesendoderm.** A maximum projection of a single cell in a Tg(*ubiP*:modErk-KTR-Clover) zebrafish embryo injected with 25 pg H2B-mScarlet-I and imaged every 1 min from ~ 4.6 hpf for 1 hr.

**Video S9. Mitotic erasure of Erk activity in the *Drosophila* anterior blastoderm.** A single z-slice through the centre of a cc13 stage nosP-modERK-KTR-T2A-His2Av-mCherry transgenic *Drosophila* embryo imaged every 3 min through to cc14.

**Video S10. Movement of zebrafish blastoderm cells at the margin.** Embryos were injected with His2B-mCherry at one-cell stage and a lateral view of the margin was imaged from 30% epiboly every 1 min for 20 min. Single nuclei were tracked in 3D and total displacement from t = 0 is shown in µm.

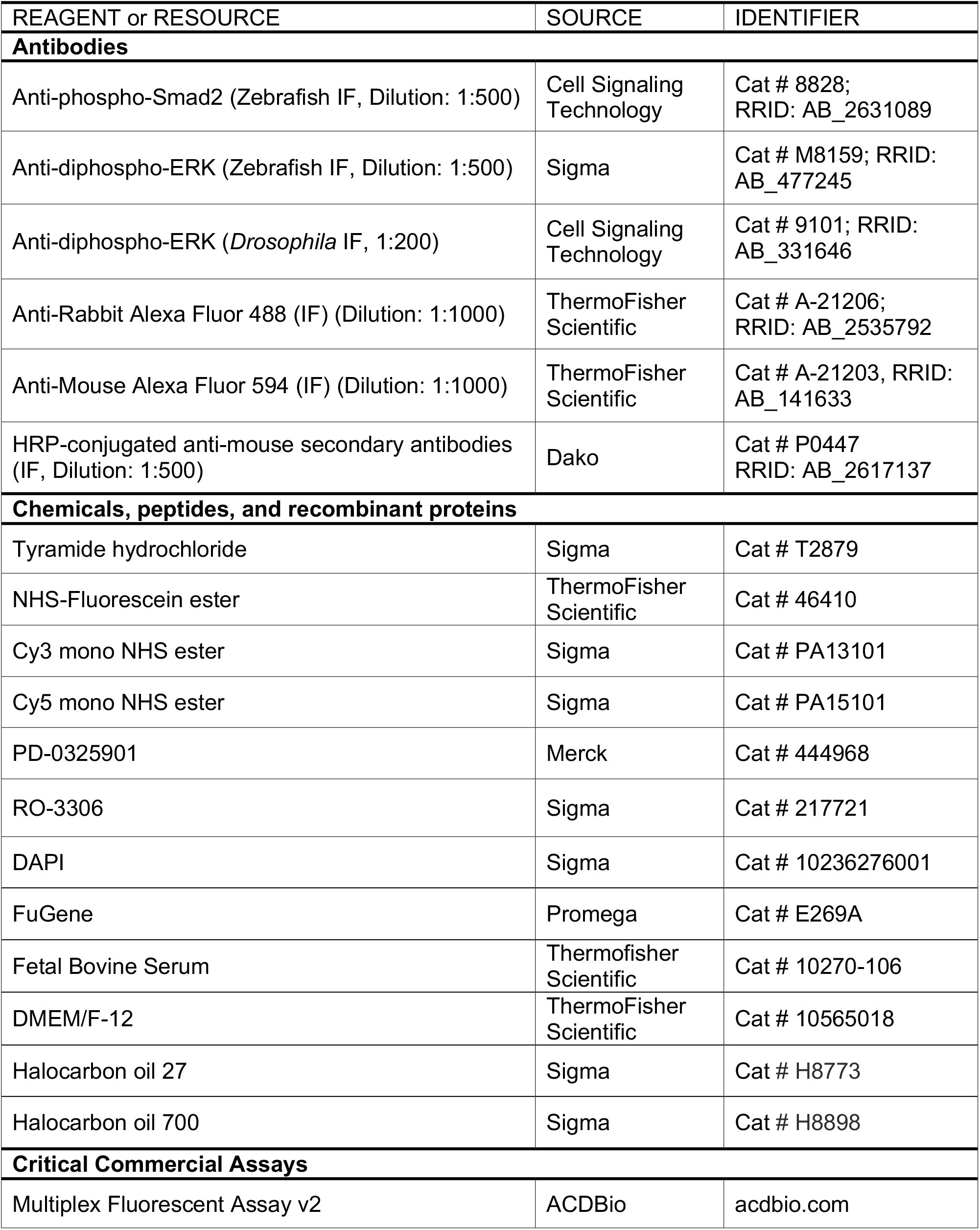
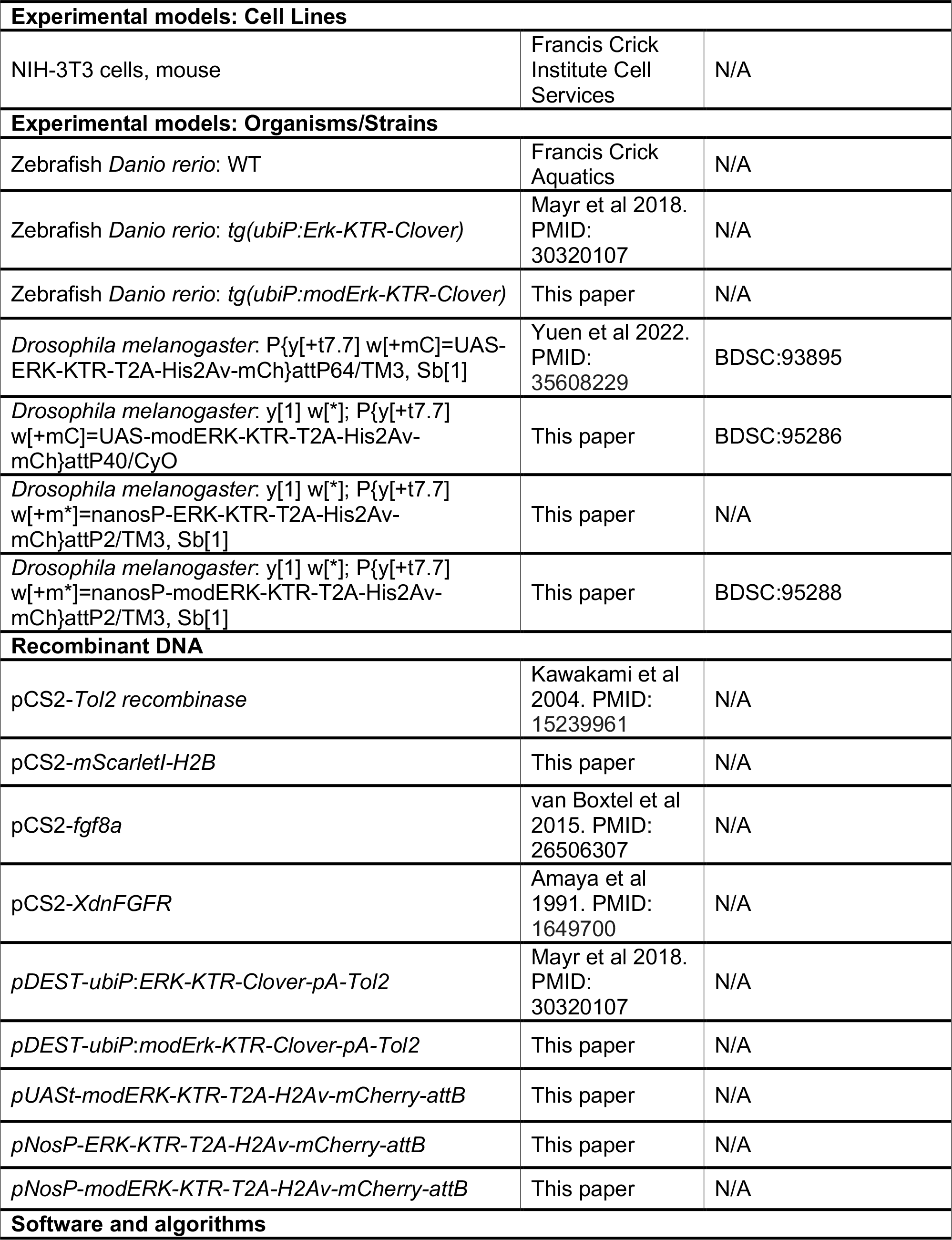
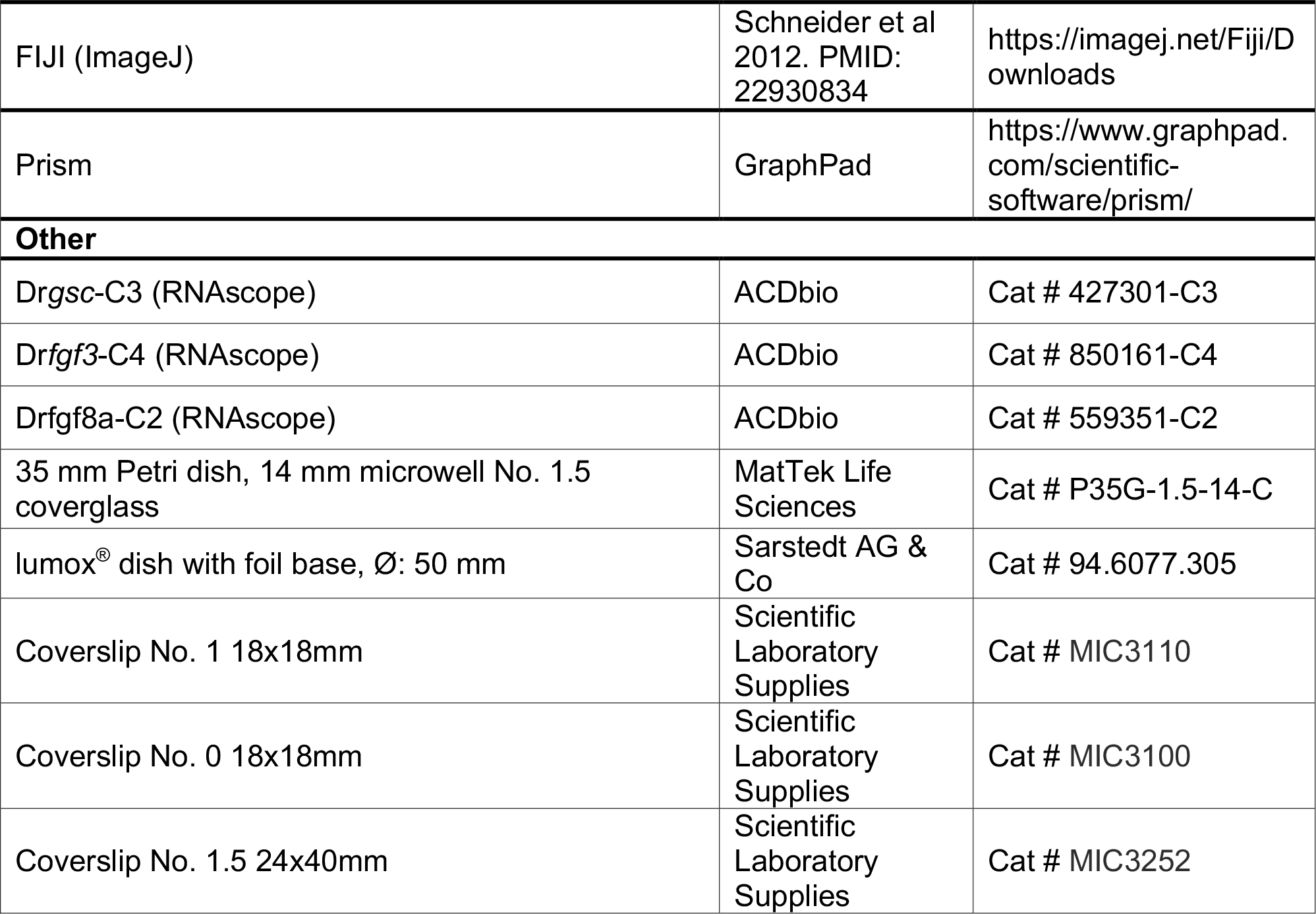
Key Resources Table

